# Dissecting PADI6 function defines oocyte cytoplasmic lattices as regulatory hubs for fundamental cellular processes

**DOI:** 10.1101/2025.02.21.639491

**Authors:** Jack P. C. Williams, Tania Auchynnikava, Afshan McCarthy, M. Teresa Bertran, Obah A. Ojarikre, Anne E. Weston, Daniel Leonce, Jessica Olsen, Kathy K. Niakan, Mark Skehel, James M. A. Turner, Louise J. Walport

## Abstract

Oocyte cytoplasmic lattices are critical for early embryo development but their composition and function are not fully understood. Mutations in *PADI6*, an essential component of cytoplasmic lattices, lead to early embryonic developmental arrest and female infertility. To investigate PADI6 function in mRNA storage, global protein levels, and lattice composition during early mammalian development we used single cell transcriptomics and proteomics methods to study two mouse models. *Padi6* null mutation resulted in inhibition of embryonic genome activation, defective maternal mRNA degradation, and disruption to protein storage on the cytoplasmic lattices. Distinct developmental phenotypes were observed with a hypomorphic *Padi6* mutation. By developing a powerful single cell proteomic fractionation method, we define the cytoplasmic lattice enriched proteome in which we find essential components of another major oocyte-specific compartment, the endolysosomal vesicular assembly (ELVA), suggesting previously unknown interconnections between them. Our findings highlight a critical scaffolding function of PADI6 and implicate cytoplasmic lattices as regulatory hubs for key processes in the oocyte and early embryo, including translation, respiration and protein degradation.

## Introduction

The growing oocyte stockpiles mRNA and proteins that are critical for successful fertilisation and early embryonic development^1–3^. The embryo relies on these maternally deposited factors because post-fertilization the embryonic genome is transcriptionally silent, until it is activated through a process called embryonic genome activation (EGA)^4–6^. These oocyte factors are important not only for EGA, but also for other key developmental processes such as the regulated degradation of maternal transcripts and proteins, the translation of early embryonic proteins, and the restructuring of cellular organelles such as the mitochondria and endoplasmic reticulum^1,7–10^. Loss or mutation of these maternal factors often results in developmental disorders and female infertility as a result of early embryonic developmental arrest^11–26^. How exactly they are stored and regulated remains unclear.

Various oocyte specific compartments and structures have been linked to the storage or regulation of different maternal factors and processes during oogenesis. Examples of such compartments include: the cytoplasmic lattice (CPL) for the storage of maternal proteins^7,27–30^, the mitochondria-associated ribonucleoprotein domain (MARDO) for the storage of maternal mRNA^31^, and the endolysosomal vesicular assembly (ELVA) for degradation of protein aggregates^32^. In each case the exact compositions are not fully characterised, with microscopy techniques having primarily been employed to determine their composition and function^7,8,30^. Various proteins are known to be essential for CPL formation, including members of the subcortical maternal complex (SCMC), comprising maternally inherited proteins such as OOEP, TLE6 and NALP5, and PADI6 (peptidyl arginine deiminase 6). Early studies suggested CPLs were composed of cytokeratin filaments, acting as a storage site of maternal ribosomes and mRNA^7,27–29,33^. Recently however, combining bulk proteomic analyses of mouse GV oocytes with super-resolution microscopy experiments and cryoelectron tomography, Schuh and co-workers reported that the CPLs are instead directly composed of the SCMC and related proteins and act as a storage hub not for ribosomes and mRNA, but for maternal proteins^30^. Association with the CPLs is proposed to be critical for the stability of these maternal factors.

PADI6 is one of the most abundant proteins in the oocyte and early embryo and is essential for the formation of the CPLs, for early embryo development and for female fertility^34–37^. More than 32 variants have been reported as the cause of infertility in studies of at least 26 women (for review see ref: ^34^). Additionally female *Padi6* knock-out mice are infertile, with embryonic arrest at the 2-cell stage^37^. PADI6 belongs to the peptidyl arginine deiminase (PADI) family, which catalyse the post-translational conversion of peptidyl arginine to citrulline in numerous protein substrates^38–42^. PADI6, however, has been reported to be inactive by *in vitro* citrullination assays against standard PADI substrates, and its activity as an arginine deiminase enzyme is yet to be confirmed^43–45^. Our recent structural characterisation of human PADI6 variants highlighted that, inconsistent with a purely scaffolding function, various embryonically lethal variants do not significantly damage the folded state of PADI6^45^. Given that PADI6 might have a catalytic function, we aimed to de-couple scaffolding functions from catalytic functions. Using microscopy, single cell transcriptomics and proteomics, we characterised oocytes and embryos from *Padi6* knock-out mice and mice with an alteration in the putative catalytic site of PADI6. This revealed that catalytic activity of PADI6 is not essential for female fertility and allowed us to separate developmental phenotypes observed due to complete absence of the CPLs, from those due to a lack of an additional function of PADI6. We then developed a single cell proteomic fractionation protocol to characterise the CPL-enriched proteome, which contained key factors linked not only to the CPLs, but also the ELVA, suggesting these structures may have interlinked functions. These analyses provide a comprehensive list of CPL-associated proteins, defining the CPLs as regulatory hubs for key cellular processes, notably protein translation, degradation, and respiration.

## Results

### Establishment of PADI6 catalytic mutant and knock-out mouse lines

To de-couple scaffolding functions of PADI6, from potential catalytic functions, we designed a mouse model in which any potential catalytic activity of PADI6 would be ablated, as a comparison to a complete null mutation. PADI catalytic activity relies on a catalytic tetrad of residues centred on a key nucleophilic cysteine^46,47^. Substitution of the catalytic cysteine in PADI2 and PADI4 results in a catalytically dead enzyme. Based on recently published high resolution X-ray crystal structures of human PADI6 (hPADI6)^45,48^, along with sequence alignments with other PADI family members and PADI6 sequences from other organisms, we predicted that any catalytic function of hPADI6 would be mediated through C675^45^, which corresponds to C663 in mouse PADI6 (mPADI6, **Figure S1A-B**). If mPADI6 is catalytically active *in vivo*, substitution of C663 with alanine (mPADI6^C663A^) would therefore ablate this function by producing a catalytically dead enzyme. Any developmental phenotype observed in mice expressing mPADI6^C663A^ could therefore be attributed to a loss of catalytic activity assuming that protein stability was unaltered.

To confirm the cysteine to alanine substitution did not affect the stability or folded structure of mPADI6, both wild-type mPADI6 and mPADI6^C663A^ were expressed and purified from mammalian cell culture (**Figure S1C-L**) and subjected to biophysical characterisation. Both the mPADI6 and mPADI6^C663A^ melted at similar temperatures and formed dimers in solution with similar ratios of monomer to dimer (**Figure S1M-P**). Dimerisation is a property of PADIs 2 to 4 and hPADI6 that has not previously been characterised for mPADI6^45–47,49^. These data indicate the C663A substitution does not affect the folded dimeric structure of mPADI6.

The *Padi6^C663A^* transgenic mouse line was then generated using CRISPR-Cas9 technology and a single-stranded oligodeoxynucleotide repair template (**Figure S2A-B**). In parallel, the C57BL/6N *Padi6^tm1a(KOMP)Wtsl^*/MbpMmucd line (ID:48953) was obtained from Mutant Mouse Resource & Research Centers and full knock-out *Padi6^-/-^* mice were generated by crossing homozygous *Padi6^tm1a^* mice with PGK-Cre mice, resulting in the deletion of exon 4 of the *Padi6* gene (**Figure S2B**). *Padi6* transcripts were not detected in *Padi6^-/-^* ovaries, and transcript levels in wild-type and *Padi6^C663A/C663A^* ovaries were not significantly different (**Figure S2C**).

### Putative PADI6 catalytic activity is not essential for female fertility

We first assessed the fertility of *Padi6^C663A/C663A^*and *Padi6^-/-^* females compared to wild-type females through natural matings with wild-type C57BL/6 males over a period of 4-5 weeks. Consistent with literature reports, *Padi6^-/-^* females were infertile, producing no litters (**Figure 1A**). *Padi6^C663A/C663A^*females on the other hand produced litters of comparable size to wild-type females, indicating that a catalytic function of PADI6 is not essential for female fertility.

**Figure 1.**
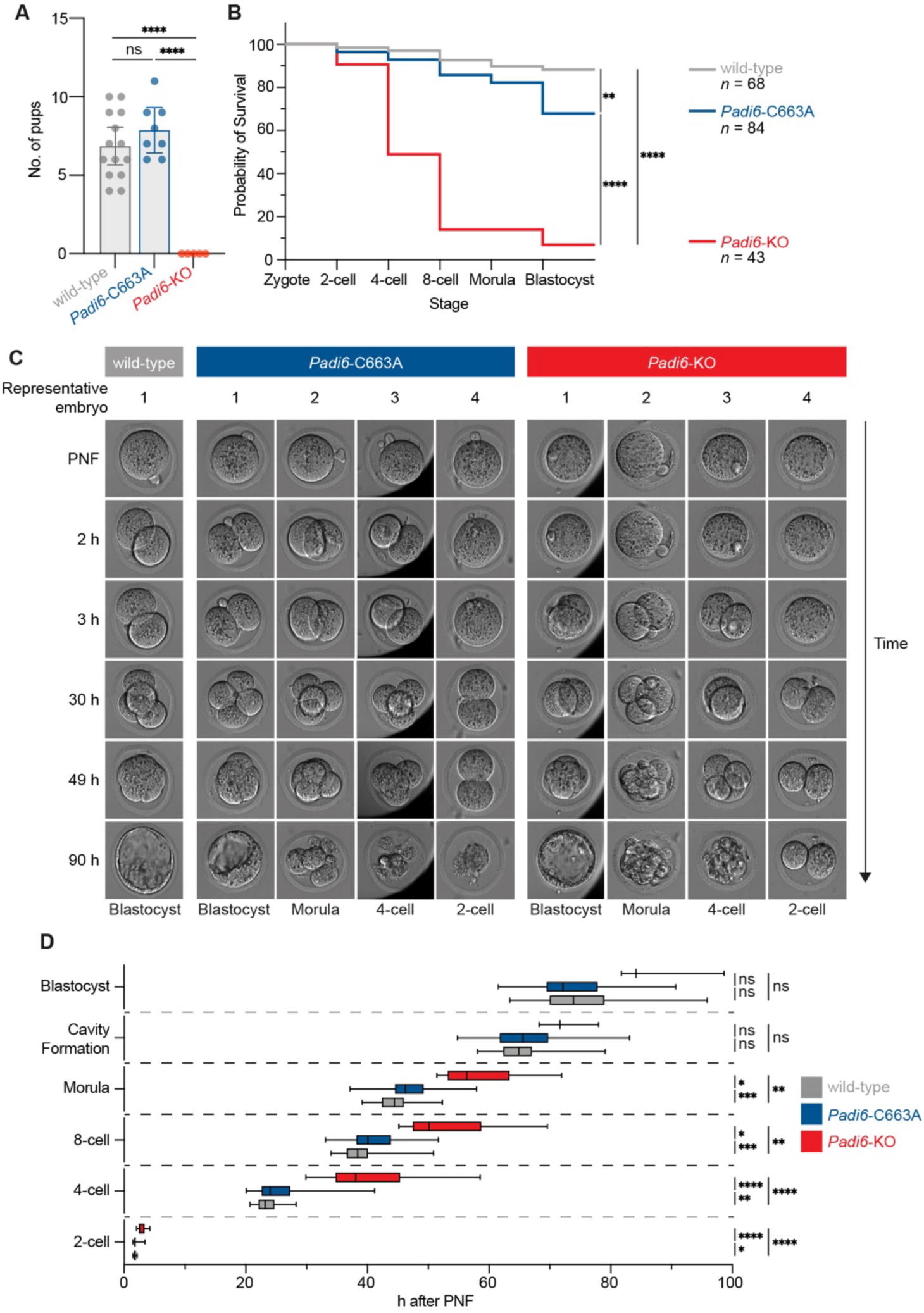
Mat-*Padi6^C663A/C663A^* embryos are partially inhibited in their development. (A) Litter size of wild-type (*n* = 14), *Padi6^C663A/C663A^*(*Padi6*-C663A, *n* = 8) and *Padi6^-/-^* (*Padi6*-KO, *n* = 5) females when naturally mated with wild-type C57BL males. Unpaired two-tailed Student’s t-test; \**P* ≤ 0.05, \*\**P* ≤ 0.01, \*\*\**P* ≤ 0.001, \*\*\*\**P* ≤ 0.0001, ns (*P* > 0.05). (B) Kaplan-Meier survival curve of wild-type (grey, *n* = 68), *Padi6*-C663A (blue, *n* = 84) and *Padi6*-KO (red, *n* = 43) embryos. Log-rank Mantel-Cox test; \**P* ≤ 0.05, \*\**P* ≤ 0.01, \*\*\**P* ≤ 0.001, \*\*\*\**P* ≤ 0.0001, ns (*P* > 0.05). (C) Representative time-lapse snapshots of embryos from wild-type, *Padi6*-C663A and *Padi6*-KO females at PNF and 2, 3, 30, 49, and 90 h post PNF. The developmental stage each embryo reached before arresting, degenerating or the end of the experiment is indicated underneath. (D) PNF normalised times at which wild-type (grey), *Padi6*-C663A (blue) and *Padi6*-KO (red) embryos reach key developmental stages. Unpaired two-tailed Student’s t-test with Welch’s correction; \**P* ≤ 0.05, \*\**P* ≤ 0.01, \*\*\**P* ≤ 0.001, \*\*\*\**P* ≤ 0.0001, ns (*P* > 0.05). The black central line of each box denotes the median value, the box represents the 25^th^ to 75^th^ percentile, and whiskers mark minimum and maximum value. PNF = Pronuclear fading.

To gain a more detailed understanding of how embryos from each line were developing, we characterised the developmental capacity of embryos from wild-type, *Padi6^C663A/C663A^* and *Padi6^-/-^*females by time-lapse imaging (**Figure 1B-C**). Unexpectedly, embryos from *Padi6^C663A/C663A^*females (mat-*Padi6^C663A/^*^C663A^) showed a small, but significant, reduction in their developmental potential compared to embryos from wild-type females. Overall, 88% of wild-type zygotes successfully reached the blastocyst stage compared with only 68% of mat-*Padi6^C663A/C663A^*embryos. 25% of mat-*Padi6^C663A/^*^C663A^ embryos arrested between the 4-cell and morula stages, with 7% failing at the zygote and 2-cell stages, compared to wild-type embryos where only 9% arrested between the 4-cell and morula stage, and 3% at zygote or 2-cell stage. This phenomenon (significant decrease in early embryo numbers in one mouse line over another despite litter sizes remaining comparable) has been observed previously and is attributed to mice, and other species, producing greater numbers of eggs and pre-implantation embryos than the uterus can accommodate, with excess embryos lost during the pre-implantation stage ^50–52^. Mat-*Padi6^C663A/C663A^* embryos were also delayed in reaching the 2-cell, 4-cell, 8-cell and morula stages (**Figure 1D**). The most significant delay was between the 4-cell and morula stages, aligning with the higher incidence of arrest observed in mat-*Padi6^C663A/C663A^* embryos occurring between these stages (**Figure 1B**). Together, these observations show an important but non-essential function of PADI6 was perturbed by the C663A substitution.

Embryos from *Padi6^-/-^* females (mat-*Padi6^-/-^*) were considerably more restricted in their development compared to both wild-type and mat-*Padi6^C663A/C663A^* embryos, with 77% arresting at the 2-cell stage, or dividing to 4-cells before undergoing fragmentation and degeneration. Six embryos (14%) did progress past the 4-cell stage, however all but three exhibited fragmentation, un-synchronised and asymmetric divisions, and eventual degeneration. These final three mat-*Padi6^-/-^*zygotes (7%) developed to blastocysts. Mat-*Padi6^-/-^* embryos were delayed in reaching every developmental stage compared to wild-type embryos, with significant delays in reaching the 2-cell to morula stages, indicative of the developmental inhibition observed when PADI6 is entirely absent.

### Cytoplasmic lattices are present in *Padi6^C663A/C663A^* germinal vesicle oocytes

Given the importance of PADI6 in CPL formation we used transmission electron microscopy to investigate whether CPLs were perturbed in *Padi6^C663A/C663A^* germinal vesicle (GV) oocytes (**Figure 2, Figure S3**). In line with previous studies, there was a complete absence of CPLs in all of the *Padi6^-/-^*GV oocytes tested^37^. Contrastingly, the CPLs were present and appeared with similar morphology and abundance in 5 out of 6 *Padi6^C663A/C663A^* GV oocytes, and 3 out of 4 wild-type oocytes tested. This confirmed that the C663A substitution in mPADI6 does not affect the capacity of PADI6 to contribute to CPL formation in mouse oocytes.

**Figure 2.**
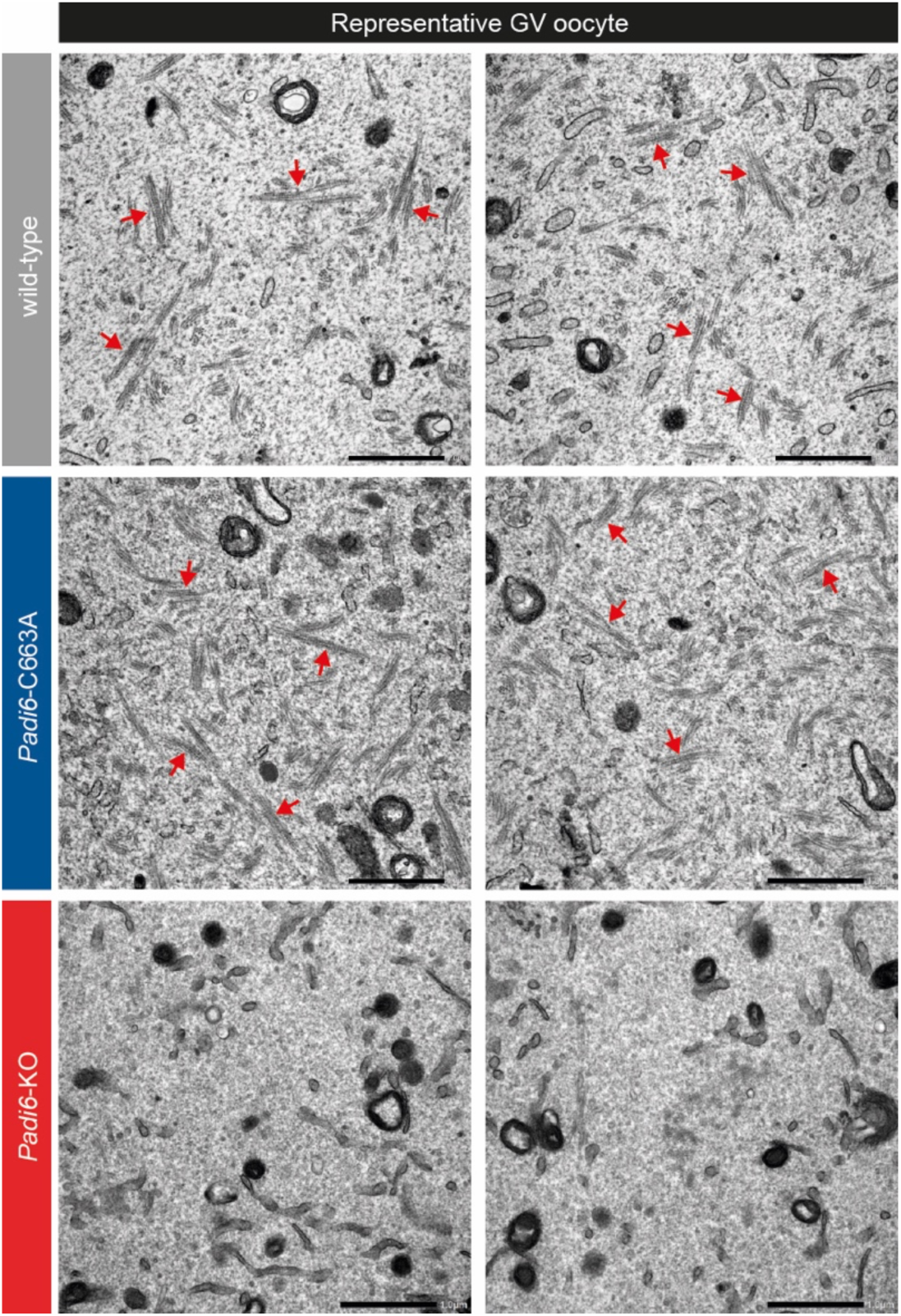
Cytoplasmic lattices are present in *Padi6^C663A/C663A^* germinal vesicle oocytes. Two representative transmission electron microscopy images from one wild-type, *Padi6^C663A/C663A^* (*Padi6*-C663A) and *Padi6^-/-^*(*Padi6*-KO) germinal vesicle (GV) oocyte showing CPLs are present in oocytes from wild-type and *Padi6*-C663A females, but not *Padi6*-KO females. Representative CPLs are indicated by red arrows. Scale bar = 1 µm. See also Figure S3.

### Transcriptome dysregulation is inversely correlated in mat-*Padi6^C663A/C663A^* and mat-*Padi6^-/-^* 2-cell embryos

To investigate the role of *Padi6* on transcriptome regulation we performed scRNA-seq on MII oocytes, zygotes, and dissociated blastomeres from 2-cell embryos (corresponding to the timing of EGA in mice) from wild-type, *Padi6^C663A/C663A^* or *Padi6^-/-^* females (**Figure 3A**). After filtering, principal component analysis (PCA) of all samples showed that cells predominantly clustered by developmental stage (**Figure S4A-D**). No reads for any other PADIs were detected in any of the samples analysed demonstrating that PADI6 is the only PADI present during these early stages, and mat-*Padi6^-/-^* oocytes and early embryos do not experience compensatory effects (**Figure S4E**). When compared to wild-type samples, both mat-*Padi6^C663A/C663A^* and mat-*Padi6^-/-^* samples displayed significant transcriptomic dysregulation. Coinciding with EGA, this dysregulation was most prominent in 2-cell embryos, where thousands of transcripts were both up and down regulated (**Figure 3B**). Unexpectedly, mat-*Padi6^C663A/C663A^* and mat-*Padi6^-/-^* embryos resulted in distinct transcriptome defects. There was little overlap between the up and downregulated genes in samples from *Padi6^C663A/C663A^* and *Padi6^-/-^* females at each stage (**Figure S4F**). Instead, an inverse correlation was observed at the 2-cell stage; 808 (58%) of the genes upregulated in mat-*Padi6^C663A/C663A^* 2-cell embryos were downregulated in mat-*Padi6^-/-^* 2-cell embryos (**Figure 3C**). This inverse effect was less prominent in the opposite direction; only 134 (18%) of the genes downregulated in mat-*Padi6^C663A^*^/*C663A*^ 2-cell embryos were upregulated in mat-*Padi6^-/-^* 2-cell embryos. Biological process gene ontology (GO) analysis of either the genes upregulated in mat-*Padi6^C663A/C663A^*, or the genes downregulated in mat-*Padi6^-/-^* 2-cell embryos, was enriched for primarily cytoplasmic translation, rRNA processing, and translation, as well as mRNA processing, RNA splicing, ribosome biogenesis, and ribosomal subunit biogenesis (**Figure S4G**). No significant enrichment for any biological processes was seen in GO analysis of either the genes downregulated in mat-*Padi6^C663A/C663A^* 2-cell embryos or upregulated in mat-*Padi6^-/-^* 2-cell embryos. This clear phenotypic divergence observed at the 2-cell stage is likely due to unique deficiencies during EGA caused by perturbation of different PADI6 functions.

**Figure 3.**
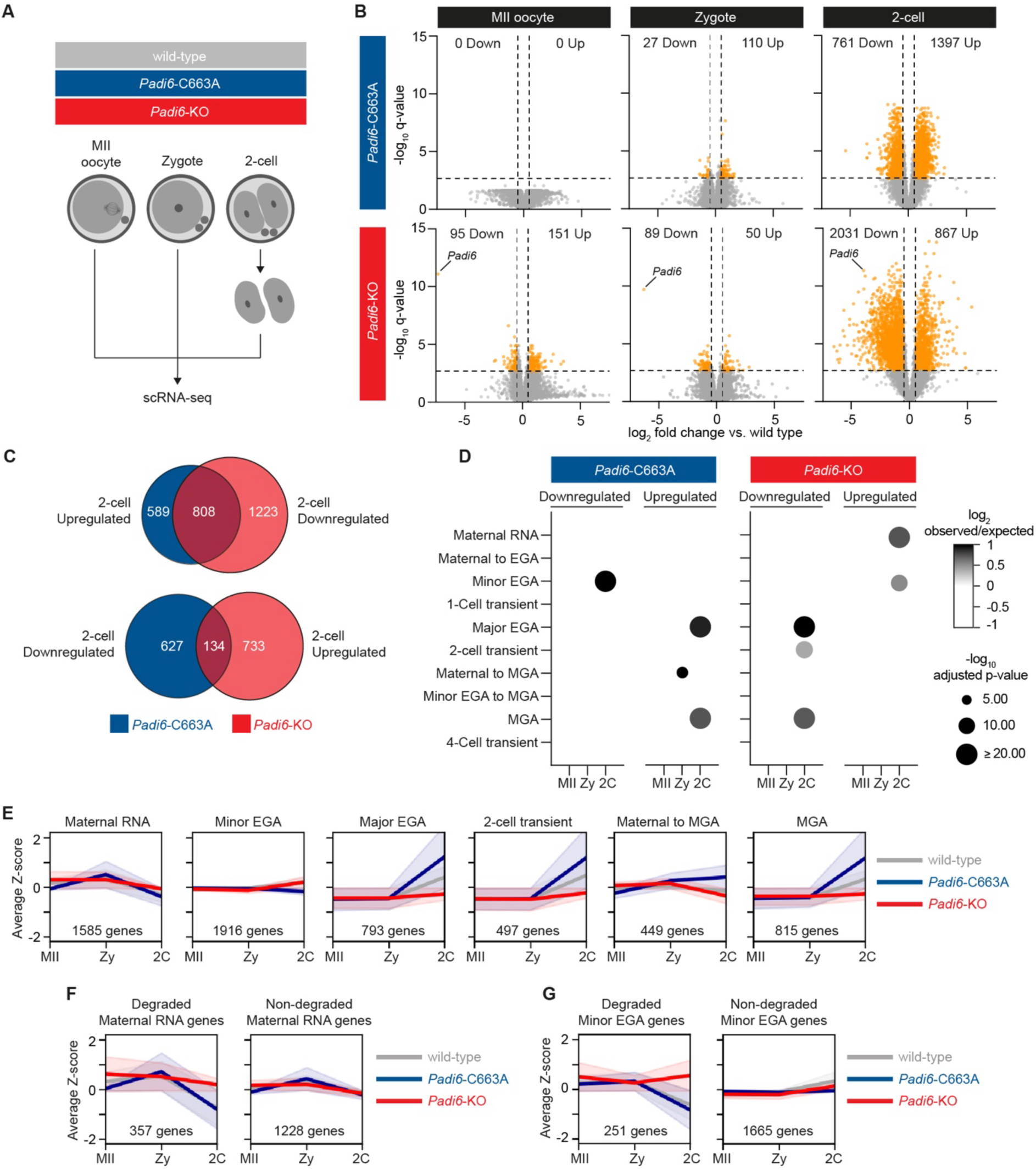
The transcriptome is dysregulated in mat-*Padi6^C663A/C663A^* and mat-*Padi6^-/-^* 2-cell embryos. (A) Schematic representation of the scRNA-seq experiment design. Per genetic background MII oocytes (*n* = 6), zygotes (*n* = 6), 2-cell blastomeres (*n* = 12). (B) Volcano plots of differentially expressed genes in mat-*Padi6^C663A/C663A^* (*Padi6*-C663A) and mat-*Padi6^-/-^* (*Padi6*-KO) MII oocytes, zygotes and 2-cell embryos compared with wild-type. Orange points represent downregulated and upregulated genes present with log2 fold change < -0.5 or > 0.5 respectively, and q-value < 0.05. Grey dashed lines indicates q-value = 0.05 and log2 fold change = -0.5 and 0.5. (C) Venn diagram of common *Padi6*-C663A 2-cell upregulated and *Padi6*-KO 2-cell downregulated genes (upper diagram) and *Padi6*-C663A 2-cell downregulated and *Padi6*-KO 2-cell upregulated genes (lower diagram). (D) Overlap of differentially expressed genes identified in (B) with DBTMEE v2 transcriptome categories^53^. Bubble plot sizes show -log10 adjusted p-values determined using a hypergeometric test for over-representation corrected for multiple testing. Bubble shade shows log2 ratio of observed gene expression changes compared with randomly expected frequencies. Under-represented genes not shown for clarity. EGA = embryonic genome activation, MGA = mid-pre-implantation gene activation. (E) Average Z-score of all genes belonging to key DBTMEE v2 transcriptome categories^53^ in MII oocytes, zygotes and 2-cell embryos from wild-type, *Padi6*-C663A and *Padi6*-KO females. Shaded area corresponds to the 95% confidence interval. (F and G) Average Z-score of all genes belonging to the maternal RNA and Minor EGA DBTMEE v2 transcriptome categories^53^ in MII oocytes, zygotes and 2-cell embryos from wild-type, *Padi6*-C663A and *Padi6*-KO females. Shaded area corresponds to 95% confidence interval. Genes are split into those that are upregulated in *Padi6*-KO 2-cell embryos and those that are not. See also Figure S4.

### EGA is dysregulated in mat-*Padi6^C663A/C663A^* embryos and inhibited in mat-*Padi6^-/-^* embryos

Differentially expressed genes were subjected to over representation analysis (ORA) by comparison with a previously defined transcriptomic database of mouse early embryo development (DBTMEE v2)^53^. Genes involved in major EGA, and mid-pre-implantation gene activation were downregulated in mat-*Padi6^-/-^* 2-cell embryos suggesting the observed developmental failure is indeed due to defective activation of the embryonic genome in the absence of PADI6. Conversely, genes upregulated in mat-*Padi6^C663A/C663A^* 2-cell embryos were linked with major EGA and mid-preimplantation gene activation (MGA), as well as 2-cell transient, and maternal to MGA genes (**Figure 3D**). Furthermore, genes belonging to the minor EGA subset were downregulated in 2-cell embryos. Together these observations highlight a possible dysregulation of EGA in *Padi6^C663A/C663A^* 2-cell embryo.

To better understand this observed dysregulation, we plotted the average Z-score of all genes in our data belonging to the maternal RNA, minor EGA, major EGA, 2-cell transient, maternal to MGA, and MGA subsets in each of our conditions (**Figure 3E**). As expected, genes belonging to subsets associated with genome activation and the 2-cell embryo (Major EGA, 2-cell transient and MGA) had increasing Z-scores between the zygote and the 2-cell stage in wild-type embryos. This increase was larger in mat-*Padi6^C663A/C663A^* embryos, further showing there were aberrantly increased transcript levels for 2-cell stage genes relative to wild-type embryos. Conversely, there was no increase in average Z-score between the zygote and the 2-cell embryo stage of mat-*Padi6^-/-^* embryos for Major EGA, 2-cell transient, and MGA genes. This confirms that PADI6 is critical for successful EGA, in line with previous reports^54,55^.

### Maternal RNA and minor EGA mRNA degradation is defective in mat-*Padi6^-/-^* 2-cell embryos

Maternal transcripts are subject to tightly regulated and specific degradation post-fertilisation^56^. Surprisingly, maternal RNA appeared upregulated in mat-*Padi6^-/-^*2-cell embryos when compared to wild-type embryos (**Figure 3D**). It has previously been reported that PADI6 is critical for the degradation of maternal RNA^54,55^. Closer inspection showed a subset of maternal RNA decreased in z-score between the zygote and 2-cell stages of wild-type and mat-*Padi6^C663A/C663A^* embryos, corresponding to their degradation (**Figure 3F**). Such a decrease was not observed in mat-*Padi6^-/-^* embryos, suggesting that the apparent upregulation of maternal RNA genes in mat-*Padi6^-/-^* 2-cell embryos is actually a result of their stabilisation or defective degradation in the absence of PADI6. A similar phenomenon was observed for minor EGA genes which were degraded between the zygote and 2-cell stage of wild-type embryos (**Figure 3G**). Together this confirms that PADI6 is essential for the regulated degradation of maternal transcripts post-fertilization.

### Establishment of a single embryo proteomic workflow

Our transcriptomic analyses revealed distinct defects in mat-*Padi6^C663A/C663A^* and mat-*Padi6^-/-^* embryos. As oocytes store proteins as well as mRNA for future development, interrogation of both the transcriptome and proteome is essential to fully understand the developmental phenotypes and functions of PADI6. Bulk analyses of *Padi6* knock-out mouse GV oocytes have recently revealed that loss of PADI6 leads to proteome defects that are decoupled from transcriptomic differences^30^. How this proteomic dysregulation extends post-fertilisation, where we observed the most dramatic developmental defects, has not yet been characterised. To address this we developed a single oocyte/embryo proteomic workflow and used it to analyse GV oocytes and 2-cell embryos from wild-type, *Padi6^C663A/C663A^* and *Padi6^-/-^* females (**Figure 4A**). The ability to profile single cells allowed us to additionally assay individual sample heterogeneity present in wild-type, *Padi6^C663A/C663A^* and *Padi6^-/-^*samples. The number of proteins identified per single cell sample in our workflow was highly consistent, with an average 5300 proteins per single GV oocyte, and 5800 proteins per single 2-cell embryo (**Figure 4B**), comparable in depth to the recently published One-Tip workflow^57^, an alternative protocol for performing single oocyte proteomics. The obtained GV oocyte proteomic data was highly consistent between samples of the same genotype (**Figure 4C**). Proteomic data obtained from 2-cell embryos showed greater variability, in particular for mat-*Padi6^C663A/C663A^* and mat-*Padi6^-/-^*embryos, likely representative of the onset of developmental disruption as observed in the time-lapse analysis (**Figure 2**).

**Figure 4.**
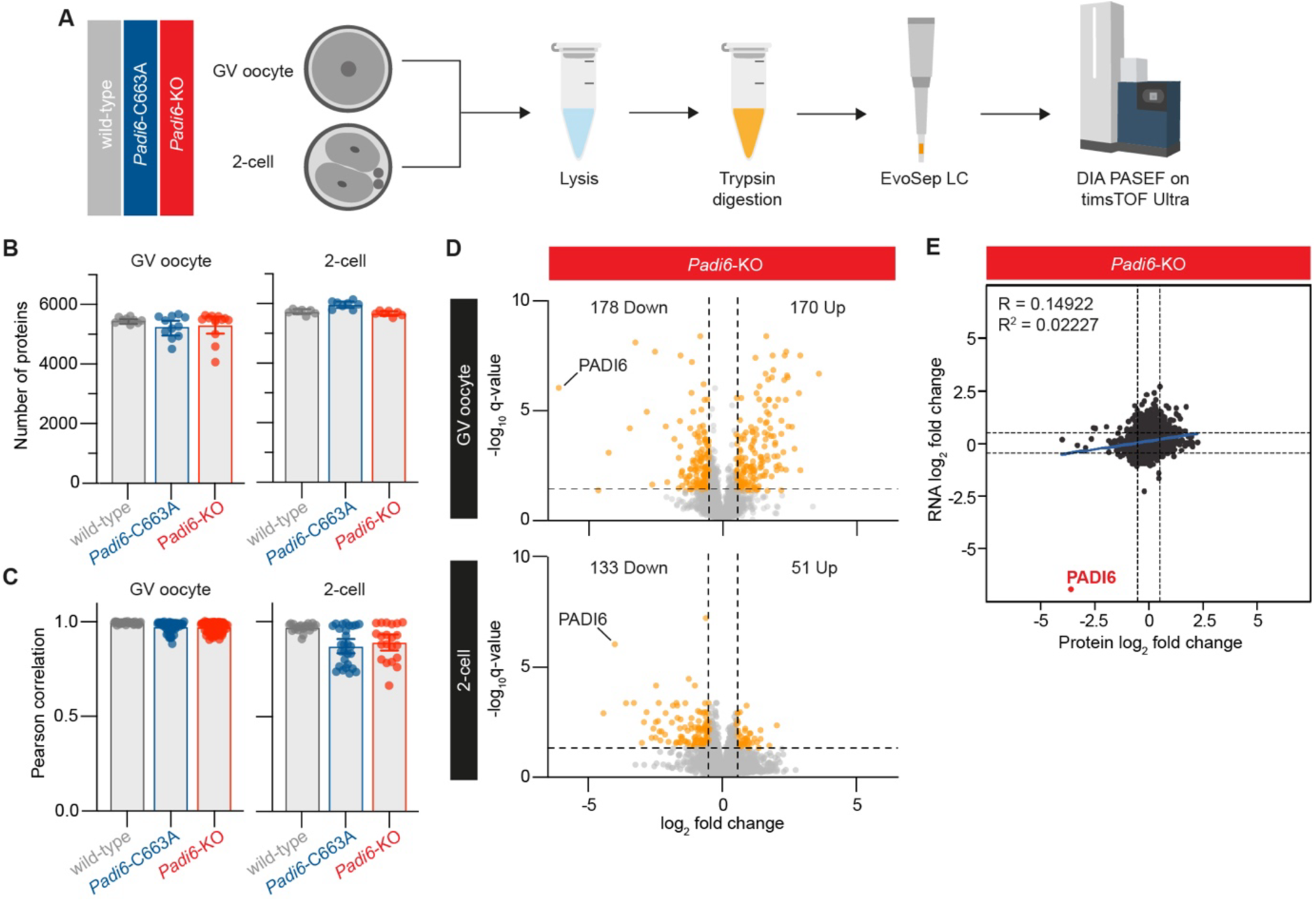
An absence of PADI6 results in significant protein dysregulation in GV oocytes and 2-cell embryos. (A) Schematic representation of the proteomic workflow used to characterise GV oocytes and 2-cell embryos from *Padi6^C663A/C663A^*(*Padi6*-C663A) and *Padi6^-/-^* (*Padi6*-KO) females. LC = liquid chromatography; DIA PASEF = data-independent acquisition parallel accumulation-serial fragmentation. (B) Number of proteins identified per single oocyte/embryo for wild-type (GV oocyte *n* = 9, 2-cell *n* = 7), *Padi6*-C663A (GV oocyte *n* = 11, 2-cell *n* = 8), and *Padi6*-KO (GV oocyte *n* = 13, 2-cell *n* = 7) GV oocytes and 2-cell embryos. (C) Pearson correlation values for wild-type, *Padi6*-C663A and *Padi6*-KO GV oocytes compared with all other GV oocytes of the same genotype. (D) Volcano plot of differentially abundant proteins in GV oocytes and 2-cell embryos from *Padi6*-KO females compared with those from wild-type females. Orange points represent downregulated and upregulated proteins present with log2 fold change < -0.5 or > 0.5 respectively, and q-value < 0.05. Dashed lines indicates q-value = 0.05 and log2 fold change = -0.5 and 0.5. (E) Protein log2 fold changes versus wild-type plotted against the log2 fold-change of their corresponding transcripts in 2-cell embryos from *Padi6*-KO females. Pearson correlation coefficient R and R^2^ values after linear regression are noted, and the fitted line plotted in blue. Dashed lines indicate q-value protein and mRNA log2 fold change = -0.5 and 0.5. See also Figure S5.

### *Padi6^-/-^* GV oocytes and 2-cell embryos show dramatic protein dysregulation

PADI6 was the second most abundant protein overall in wild-type GV oocytes and the 13^th^ most abundant in wild-type 2-cell embryos (**Figure S5A-B**), consistent with bulk proteomic data^35^. PADI6 protein level in both GV oocytes and 2-cell embryos from *Padi6^C663A/C663A^* females was not significantly different to wild-type levels, in line with our biophysical data showing the C663A substitution did not affect protein expression or stability (**Figure S5C-D**).

Globally, in *Padi6^C663A/C663A^* GV oocytes no proteins appeared dysregulated when compared to wild-type (**Figure S5E**). Dramatic dysregulation was observed in *Padi6^-/-^* GV oocytes with 178 proteins downregulated and 170 proteins upregulated (**Figure 4D**). At the 2-cell stage, in contrast to the dramatic transcriptional dysregulation observed in mat-*Padi6^C663A/C663A^* embryos (**Figure 3B**), no proteins were significantly dysregulated (**Figure S5E**). Again, as in the GV oocyte, protein dysregulation was seen in mat-*Padi6^-/-^* 2-cell embryos with 133 downregulated and 51 upregulated (**Figure 4D**), although this is still less substantial than the transcriptomic dysregulation observed in equivalent embryos (**Figure 3B**). Overall, little to no correlation was observed between mRNA and protein dysregulation in mat-*Padi6^-/-^*2-cell embryos (**Figure 4E**). This is consistent with data showing that minimal translation occurs in the 2-cell embryo, with the first large translational wave only occuring in the morula^58^, suggesting that transcriptomic and proteomic dysregulation need not be linked at early developmental stages.

Pathway enrichment analysis of the dysregulated proteins using the Search Tool for the Retrieval of Interacting Genes/Proteins (STRING)^59^ showed no significant enrichment for any biological processes for those downregulated in the *Padi6^-/-^*GV oocytes. On the other hand, numerous metabolic and catabolic pathways were upregulated in *Padi6^-/-^* GV oocytes, indicating energy production pathways may be altered in the absence of PADI6 (**Figure S5F**). Mat-*Padi6^-/-^* 2-cell downregulated proteins were not significantly enriched for any biological processes, however, mat-*Padi6^-/-^* 2-cell upregulated proteins enriched significantly for translation, and mRNA processing and splicing processes, suggesting that early genome activation events could be misregulated due to defects in RNA processing and translation (**Figure S5G**).

The levels of key CPL scaffolding proteins from the SCMC (OOEP, TLE6, NALP5 and KHDC3), were not significantly different in either *Padi6^C663A/C663A^*or *Padi6^-/-^* GV oocytes or 2-cell embryos when compared to wild-type (**Figure S5H-I**). UHRF1 was abundantly expressed in wild-type GV oocytes and 2-cell embryos, but virtually absent in *Padi6^-/-^* GV oocytes and 2-cell embryos (**Figure S5H-I**). This PADI6-dependent co-depletion of UHRF1 on the protein level was previously observed in bulk proteome analyses^30^, and is not mirrored by a corresponding decrease in the mRNA level of UHRF1 (**Figure S5J**), suggesting PADI6 regulates UHRF1 at the protein level. No significant change in UHRF1 level was observed in GV oocytes or 2-cell embryos from *Padi6^C663A/C663A^* females compared with wild-type, ruling out a catalytic contribution to this stabilisation.

### The CPLs store critical housekeeping proteins for the early embryo

Dysregulation and developmental defects in *Padi6^-/-^* GV oocytes have been linked to the absence of CPLs, which are proposed to be key regulators of protein stability in mammalian oocytes^30^. Multiple studies have highlighted the complex composition of CPLs, which appear to house multiple proteins in addition to the core members. Key structural components of the CPLs have been identified (PADI6, SCMC, UHRF1)^37,60–63^, along with the recent identification of a range of proteins stored on the lattices^30^. Despite this growing understanding, however, the full composition remains uncharacterised. A complete list of the CPL components would aid in our understanding of CPL function and the phenotypes related to PADI6.

CPLs are Triton X-100 insoluble and historically triton extraction experiments coupled with immunofluorescence have been employed to characterise CPL-associated factors^7,8,33^. We reasoned that Triton X-100 treatment followed by single oocyte mass spectrometry of *Padi6^-/-^*and wild-type oocytes could be used to more comprehensively identify proteins associated with intact CPLs. CPL associated proteins would be enriched in the Triton X-100 insoluble fraction in wild-type oocytes after treatment, but washed out in *Padi6^-/-^* oocytes that lack the CPLs (**Figure 5A**). Wild-type, *Padi6^C663A/C663A^*and *Padi6^-/-^* GV oocytes were subjected to Triton X-100 treatment immediately after harvesting, washed thoroughly in PBS and analysed using our single cell proteomics workflow. As expected for sub-cell proteomics, significantly fewer proteins per oocyte were identified in Triton X-100 treated samples, with an average of 1700 (**Figure 5B**). Overall sample variation for treated samples increased in comparison with intact oocytes; as seen previously, more variation was also observed in *Padi6^C663A/C663A^* and *Padi6^-/-^* GV oocytes than wild-type potentially indicating heterogeneity in CPL defects (**Figure 5C**).

**Figure 5.**
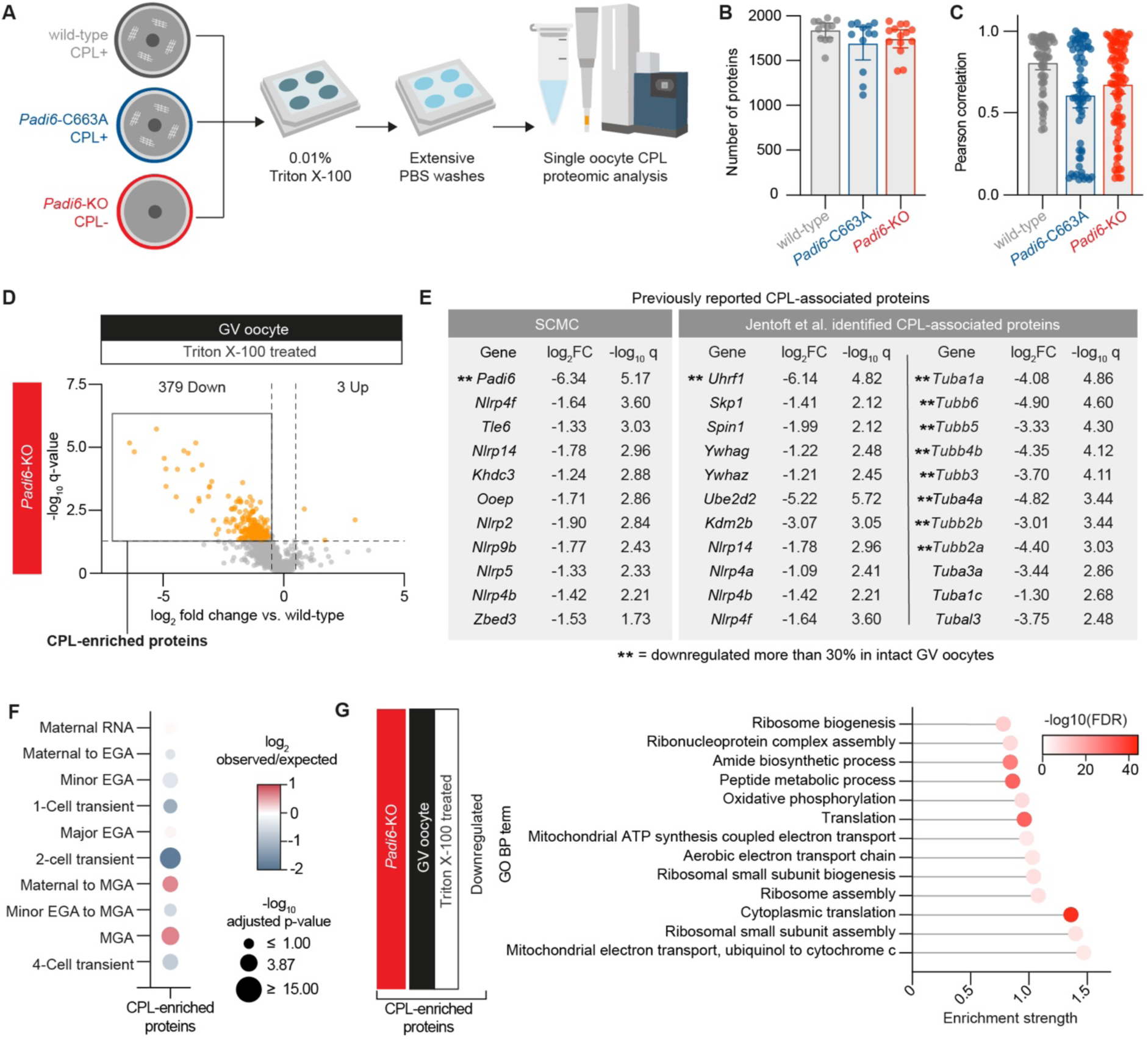
Characterisation of the CPL-enriched proteome. (A) Schematic representation of the experimental workflow used to identify CPL associated proteins in GV oocytes from *Padi6^C663A/C663A^* (*Padi6*-C663A) and *Padi6^-/-^* (*Padi6*-KO) females. (B) Number of proteins identified per single oocyte for wild-type (*n* = 12), *Padi6*-C663A (*n* = 12), and *Padi6*-KO (*n* = 14) Triton X-100 treated GV oocytes. (C) Pearson correlation values for wild-type, *Padi6*-C663A and *Padi6*-KO Triton X-100 treated GV oocytes compared with all other GV oocytes of the same genotype. (D) Volcano plot of differentially abundant proteins in Triton X-100 treated GV oocytes from *Padi6*-KO females compared with Triton X-100 treated GV oocytes from wild-type females. Orange points represent downregulated and upregulated proteins present with log2 fold change < -0.5 or > 0.5 respectively, and q-value < 0.05. Dashed lines indicates q-value = 0.05 and log2 fold change = -0.5 and 0.5. Downregulated proteins are classed as CPL-enriched. (E) Previously reported CPL-associated proteins present in the CPL-enriched fraction of (D). Proteins grouped into SCMC components and the CPL proteins identified by Jentoft et al.^30^ with log2 fold changes and -log10 q-values as per (D). ** = downregulated >30%, intact *Padi6*-KO GV oocytes compared with wild-type. (F) Overlap of identified CPL-enriched proteins with DBTMEE v2 transcriptome categories^53^. Bubble plot sizes show -log10 adjusted p-values determined using a hypergeometric test for over-representation corrected for multiple testing. Bubble colour shows log2 ratio of observed protein level changes compared with randomly expected frequencies. EGA = embryonic genome activation, MGA = mid-pre-implantation gene activation. (G) Enriched GO BP terms after STRING analysis of CPL-enriched proteins. Dot colour indicates -log10 false discovery rate (FDR). GO BP terms were filtered prior to plotting to include only those with an enrichment strength > 0.75 and FDR < 0.00001. See also Figure S6.

379 proteins were enriched in wild-type in comparison to *Padi6^-/-^*Triton X-100 treated oocytes (**Figure 5D**). Given the defect in CPL formation, we propose that most of these proteins are ones that are rendered insoluble through association with the CPLs in wild-type oocytes and hence comprise a bona fide CPL-associated proteome. Supportive of this interpretation among the enriched proteins were all reported members of the mouse SCMC (OOEP, KHDC3L, TLE6, NALP5, NLRP2, NLRP4F, NLRP7, ZBED3)^9,10,62,64–67^, as well as novel CPL proteins identified by Jentoft *et al.* using expansion microscopy (UHRF1, SKP1, SPIN1, YWHAZ, a-Tubulin, UBE2D2, KDM2B, NLRP14, NLRP4A, NLRP4B, NLRP4F)^30^ (**Figure 5E**).Together these results indicate that we have established a powerful sub-cellular proteomic workflow from single mouse oocytes capable of identifying proteins associated with the CPLs. No proteins were significantly downregulated in *Padi6^C663A/C663A^* Triton X-100 treated oocytes (**Figure S6A**), in line with our previous observations that the CPLs are intact in these oocytes (**Figure 2**).

Association of proteins with the CPLs has been proposed to promote stability and storage of proteins in the oocyte^30^. Whilst we observe more CPL-enriched proteins were downregulated than upregulated in intact *Padi6^-/-^* GV oocytes, the majority of proteins (330/379) displayed little to no change in abundance suggesting association to the CPLs is not a determinant of their stability (**Figure S6B**). Maternal to mid-preimplantation genome activation proteins were the most overrepresented subset when compared with the DBTMEE v2 database, in which 94 proteins were CPL-enriched^53^ (**Figure 5F**). These proteins include a variety of factors involved in key cellular processes such as translation, protein degradation, respiration and splicing. Due to their critical functions these proteins are uniformly expressed from the oocyte to blastocyst stage^53^. STRING analysis of the CPL-enriched proteins showed a significant enrichment for both translation, and ATP generation biological processes (**Figure 5G**). This finding suggests the CPLs are key structures in the regulation and storage of crucial housekeeping proteins during early embryo development.

### Protein translation, regulation and degradation machinery are enriched in the CPLs

Early work into the CPLs reported that the CPLs act as a storage hub for maternal ribosomes and mRNA, however this theory was recently disputed^7,30^. In our data, 28 large ribosomal subunit proteins and 19 small ribosomal subunit proteins were enriched in wild-type compared to *Padi6^-/-^* Triton X-100 treated GV oocytes (**Figure 6A**). This highlights that there is indeed a relationship between the CPLs and maternal ribosomes that is dependent on PADI6. Given the number of ribosomal subunits identified, this likely represents intact ribosomes.

**Figure 6.**
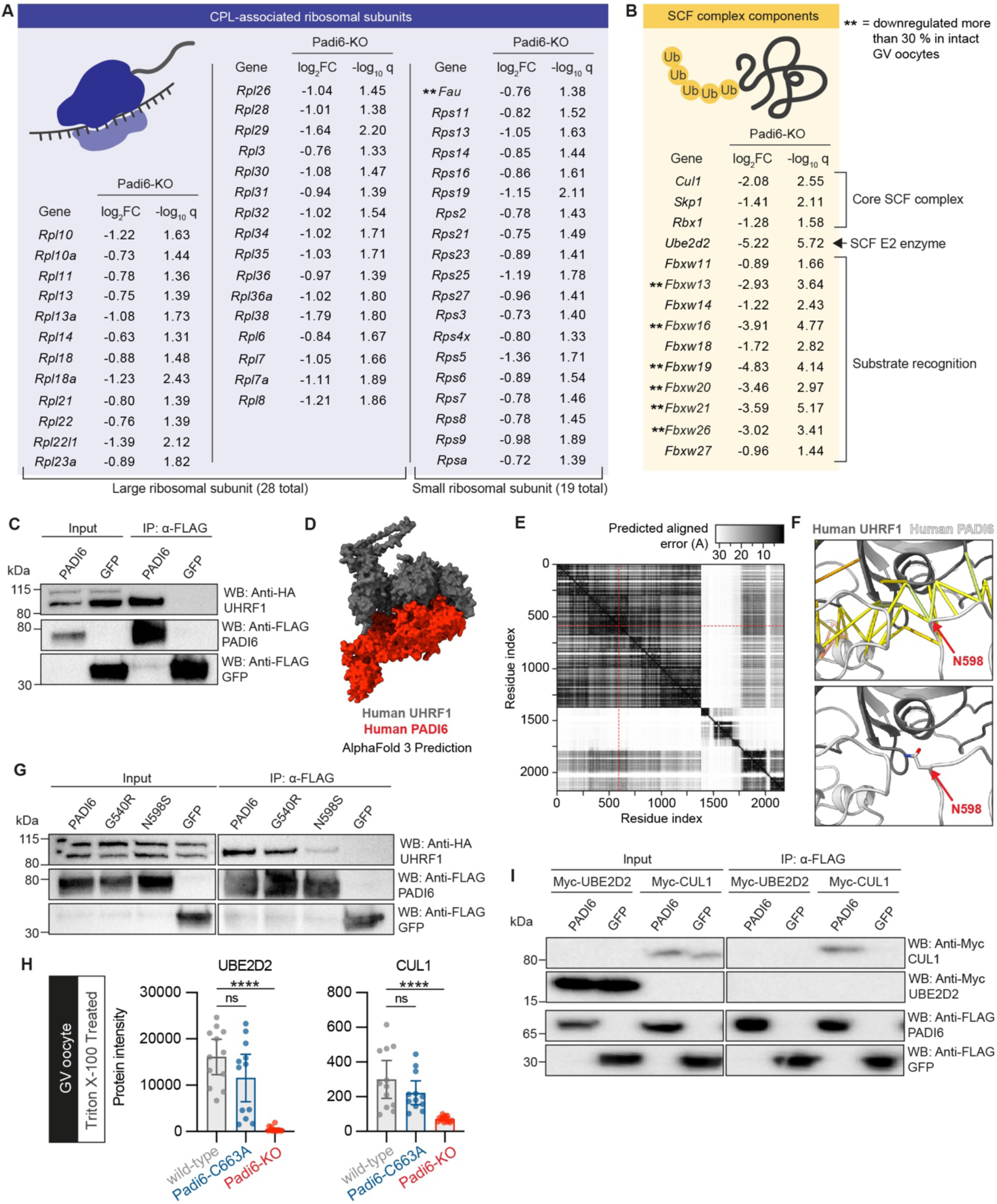
The CPLs are associated with protein translation, regulation and degradation machinery, and the oocyte mitochondria. (A) All large (28 total) and small (19 total) ribosomal sub-units found to be CPL-enriched with the log2 fold-change and -log10 q-values of Triton X-100 treated *Padi6^-/-^* (*Padi6*-KO) GV oocytes compared with Triton X-100 treated wild-type GV oocytes. ** = proteins downregulated more than 30% in intact *Padi6*-KO GV oocytes. (B) Proteins associated with the SCF complex found to be CPL-enriched with log2 fold changes and -log10 q-values as per (A). ** = proteins downregulated more than 30% in intact *Padi6*-KO GV oocytes. (C) Western blot analysis of co-immunoprecipitation experiments of HEK-293T lysates (Input) using FLAG(M2)-beads. Cells were transiently co-transfected with either FLAG-hPADI6 or FLAG-GFP and HA-UHRF1. Western blots probed with antibodies indicated in figure. Experiment performed in biological duplicate. Representative blot shown. (D) Model 0 of the AlphaFold Server^72,73^ predicted interaction between hPADI6 (red) and UHRF1 (grey). Visualised with ChimeraX^74,75^. (E) Predicted aligned error heatmap of the interaction between hPADI6 and UHRF1. hPADI6-N598 is highlighted by a red dashed line. Residue indices: hPADI6 molecule 1: 0-693, hPADI6 molecule 2: 694-1388, UHRF1 molecule: 1389-2182. (F: Upper) Close-up of N598 at the interaction interface with predicted interactions with UHRF1 shown. N598 is highlighted. Visualised with ChimeraX^74,75^. The predicted error of contacts are coloured from low error strong contacts to high error weak contacts, blue to yellow to red. (F: Lower) Close up of N598 in the AlphaFold Server^72,73^ predicted hPADI6 (white) and UHRF1 (grey) interaction surface. Model 0 is shown and N598 highlighted. Visualised with ChimeraX^74,75^. (G) Western blot analysis of co-immunoprecipitation experiments from HEK-293T lysates (Input) using FLAG(M2)-beads. HEK-293T cells were transiently co-transfected with HA-UHRF1, and either FLAG-hPADI6, FLAG-hPADI6^G540R^, FLAG-hPADI6^N598S^, or FLAG-GFP. Western blots probed with antibodies indicated in figure. Experiment performed in biological duplicate. Representative blot shown. (H) Protein intensity values of UBE2D2 and CUL1 in Triton X-100 Treated GV oocytes from wild-type (n = 12), *Padi6^C663A/C663A^*(*Padi6*-C663A, n = 12) and *Padi6^-/-^* (*Padi6-*KO, n =14) females. Unpaired two-tailed Student’s t-test; \**P* ≤ 0.05, \*\**P* ≤ 0.01, \*\*\**P* ≤ 0.001, \*\*\*\**P* ≤ 0.0001, ns (*P* > 0.05). (I) Western blot analysis of co-immunoprecipitation experiments from HEK-293T lysates (Input) using FLAG(M2)-beads. Cells were transiently co-transfected with either FLAG-hPADI6 or FLAG-GFP and Myc-UBE2D2 or Myc-CUL1. Western blots probed with antibodies indicated in figure. Experiment performed in biological duplicate. Representative blot shown. See also Figure S6 and S7.

Various proteins involved in protein degradation, including SKP1, are known to be associated with the CPLs^30^. As well as SKP1, we find both CUL1 and RBX1, the other two core components of the SKP1-CUL1-F-box protein (SCF) ubiquitin ligase complex, and UBE2D2, a known E2 of the complex, to be CPL-enriched (**Figure 6B**). The SCF complex mediates the ubiquitination of proteins involved in the cell cycle and transcription, and interacts with F-box proteins as substrate recognition adaptors ^68–71^. 10 F-box and WD-40 domain (FBXW) proteins were also present in the CPL-enriched fraction, including the well characterised SCF substrate recognition adapter FBXW11 (**Figure 6B**). These FBXW proteins could resemble an array of embryo specific targeting adaptors for the regulated and timely degradation of maternal factors. Along with this ubiquitination machinery, 7 proteosomal subunits were enriched on the CPLs (**Figure S6C)**. Proteosomes themselves are stored along with aggregated proteins, endolysosomes and autophagosomes in the liquid-like matrix ELVAs^32^. This protein matrix is formed by RUFY1 and ELVAs contain the lysosome associated LAMP1, both of which were also enriched in Triton X-100 treated wild-type oocytes compared to *Padi6* knock-out (**Figure S6D**). Together this indicates that as well as translation machinery, the CPLs house and/or regulate protein degradation machinery in the oocyte.

### The CPLs are associated with oocyte mitochondria

Intriguingly, many proteins involved in ATP generation and mitochondrial respiration, in particular components of the electron transport chain, were enriched in the CPLs (**Figure S6E**). 26 components of Respiratory Complex I, III, IV and ATP synthase machinery were CPL-enriched. This data highlights a relationship between the mitochondria and CPLs, which is dysregulated in the absence of PADI6. Indeed an absence of PADI6 in the oocyte has been shown to result in defects in mitochondrial localisation^8^. Notably, we also identify three key proteins (GPX1, SOD2, and PRDX6), which neutralise reactive oxygen species (ROS)^76^, and are upregulated in times of oxidative stress, that are upregulated in *Padi6^-/-^* GV oocytes (**Figure S6F**). This highlights ROS damage, possibly caused by mitochondrial disruption, as a contributor to the severe developmental phenotypes present in the absence of PADI6.

### Ubiquitination machinery associates with the CPLs through PADI6

Finally, with an atlas of CPL proteins in hand, we sought to validate whether association of some of the proteins we identified as CPL-associated was mediated by PADI6. Due to the high numbers of mice required to perform co-immunoprecipitation (co-IP) experiments directly from oocytes, we instead chose to validate select candidate proteins as PADI6 interactors by co-immunoprecipitation of FLAG-PADI6 and each candidate protein expressed in transiently transfected HEK-293T cells. Given the dramatic depletion of UHRF1 in *Padi6* knock-out oocytes and embryos (**Figure S5H-I**), we chose to investigate whether this was a direct interaction. We first validated by co-IP that human PADI6 (hPADI6) interacted with human UHRF1 when both were transiently transfected into HEK-293T cells (**Figure 6C**). AlphaFold 3^72,73^ modelling of a possible hPADI6 UHRF1 interaction produced five predictions with the binding surface in each centred on N598 of hPADI6 (**Figure 6D-F, Figure S7**). We have previously shown that N598 is surface exposed in hPADI6 and its substitution does not affect thermodynamic stability^45^.

Interestingly, a p.N598 *PADI6* variant (p.N598S) has been reported in an infertile woman who experienced longer pregnancies, high miscarriage incidence and hydatidiform mole formation^77^. Given the role of UHRF1 in DNA methylation and suggestions that imprinting linked methylation defects can be a contributing factor towards hydatidiform mole formation^78^, we tested whether the PADI6 interaction with UHRF1 was disrupted when N598 was substituted to S. In support of the AlphaFold prediction, substitution of N598 to S in hPADI6 dramatically reduced the capacity of hPADI6 to pull-down HA-UHRF1 (**Figure 6G**). In comparison, the surface exposed and clinically significant variant p.G540R (phenotype: early embryonic arrest^79^) pulled down HA-UHRF1 to a comparable level as the wild-type protein. These data indicate that PADI6 interacts directly with UHRF1 on the CPLs, and that this interaction is critical for the stability of UHRF1, and embryo development.

We additionally identified the SCF E3 ligase complex as CPL-enriched, with core component CUL1 being among the components enriched in wild-type compared to *Padi6^-/-^* Triton X-100 treated oocytes (**Figure 6H**), indicating significant disruption in the absence of PADI6. We were also able to validate this as a direct interaction; hPADI6 interacted with CUL1 when transiently transfected into HEK-293T cells (**Figure 6I**). It did not pull-down known SCF E2 enzyme UBE2B2, which we also found to be CPL-associated suggesting association of other members of the SCF complex may be mediated by the complex itself (**Figure 6H-I**). Together, these findings support a function of the CPLs as regulatory hubs for protein ubiquitination and degradation, with key interactions mediated through PADI6.

## Discussion

PADI6 is critical for early embryo development in mice and humans, but aside from being integral to CPL formation, the functions of PADI6 are unclear. In this work we have used two mouse models, a knock-out and a single amino acid substituted line, to untangle CPL scaffolding dependent functions, from potential catalytic functions. We further leveraged these models to develop a proteomic workflow that characterised the CPL-enriched proteome, providing the most comprehensive catalogue of their components to date. Our triton extraction proteomic workflow was able to identify all validated CPL components, as well as previously unknown factors, in a single experiment, contrasting with previous studies that have identified small numbers of CPL-associated factors through candidate-based microscopy approaches.

Our data validates previous hypotheses on the composition of the CPLs and highlights potential previously unknown functions including protein degradation. We identify the entire SCF complex, a known cooperating E2 enzyme and a battery of potential substrate targeting adaptors as CPL-enriched. Of the many candidates we identified we further characterised the interaction of two ubiquitination factors, UHRF1 and CUL1, with PADI6 indicating they are likely associated to the CPLs through direct interaction with PADI6. Although we see a bias towards downregulation of proteins associated with the CPLs in their absence, on the whole this effect is relatively subtle, suggesting that the main function of the CPLs may be the localisation and storage of proteins, rather than their stabilisation, with the stabilisation of certain proteins being a downstream consequence of their immobilisation.

Many oocyte specific super structures with unique functions have been reported in addition to CPLs^30^ including the MARDO^31^ and ELVA^32^. However, with microscopy techniques primarily being employed in their characterisation, the exact protein compositions and regulatory mechanisms of each are unknown. Additionally, the interplay between these structures has not been studied. Key factors required for the formation of these structures have been identified, notably PADI6 and the SCMC for the CPLs^37,60–63^, and RUFY1 for the ELVA^32^. RUFY1 appeared in our CPL-enriched proteome. Given the ability of RUFY1 to form phase-separated structures in the oocyte, it is conceivable that it also contributes to CPL formation and scaffolding. Or alternatively, this reveals an interdependency between the ELVA and CPLs. In support of this, we find proteasome and ubiqutination machinery enriched in the CPLs. Further work is required to untangle the interrelations between these oocyte structures, and the commonality of proteins required for their formation and regulation.

Many variants in CPL associated genes are reported to result in severe developmental phenotypes, most commonly early embryonic developmental arrest and female infertility. For the most damaging mutations, the most common observation attributed to the developmental phenotypes is an absence of the CPLs. Our work however, highlights that there is more nuance to the function of these proteins. Clinical data also supports this, with less damaging mutations in *PADI6* having been reported to result in longer pregnancies resulting in miscarriages, and in fertile women whose offspring develop multi-locus imprinting disorders. This is exemplified by the relationship between PADI6 and UHRF1. An absence of PADI6 leads to an almost complete depletion of UHRF1 in the oocyte and early embryo in this and other studies in mice alongside loss of the CPLs. However, presence of the hPADI6 p.N598S variant, which we demonstrate significantly disrupts the binding of PADI6 and UHRF1, does not result in early embryonic developmental arrest. Instead, embryos develop beyond the early stages and form molar pregnancies resulting in miscarriages. Based on our data we propose that this is a consequence of the mutation disrupting UHRF1 binding, but not preventing CPL formation. To fully understand the functions of these proteins, both in the context of the CPLs, and not, models incorporating these more subtle protein changes will be required.

We have shown that a catalytic function of PADI6 is not essential for the formation of the CPLs, or more broadly for early embryo development and female fertility. Embryos from *Padi6^C663A/C663A^* females do, however, show a significantly higher incidence of arrest compared to embryos from wild-type females, and 2-cell embryos showed dramatic transcriptomic dysregulation, in particular in major EGA factors. The possibility of a non-essential catalytic function of PADI6 in oogenesis and early embryo development cannot therefore be ruled out, potentially in the epigenetic regulation of transcription similar to PADI4. Aside from the transcriptomic and EGA dysregulation observed at the 2-cell stage, we found that a large subset of mat-*Padi6^C663A/C663A^* embryos arrested or degenerated at the morula stage. This points toward additional functions of PADI6 later on in development, which could be catalytic and cannot easily be studied in knock-out models due to the high incidence of arrest at the 2- to 4-cell stages. It is conceivable that a possible catalytic function of PADI6 in the morula could be a means to enable the dissociation of the CPLs, akin to the way in which histone citrullination by PADI4 can weaken the electrostatic interactions between histone tails resulting in the de-compaction of chromatin^46,80,81^. The CPLs have been reported to persist until the morula stage, and the mechanism of their disassembly is unknown^58^. This disappearence of the CPLs is accompanied by a large wave of embryonic translation and an increase in free ribosomes. Our data shows that ribosomes are associated with the CPLs. It would be of interest in future work to determine whether the CPLs persist beyond the morula stage in mat-*Padi6^C663A/C663A^* embryos and whether this is accompanied by inhibited translation.

### Limitations of the study

We have elucidated the composition of the CPLs in mouse oocytes, however, it has been shown that the morphology of CPLs in human oocytes is different, suggesting there may also be differences in their composition. Additionally, although we can rule out an essential role of catalysis in female fertility, due to the difficulties in confidently de novo assigning citrullination, in particular from limited sample quantity, we have not probed directly for citrullinated proteins in our samples and so cannot completely rule out its presence. The development of better tools and methods is needed to be able to study citrullination in the context of oogenesis and early embryo development. Without this, the catalytic activity of PADI6 cannot be confidently assigned.

## Resource Availability

### Lead contact

Requests for further information and resources should be directed to and will be fulfilled by the lead contact, Louise Walport.

### Materials availability

Resources generated in this study are available from the lead contact without restriction.

### Data and code availability

- Raw and processed scRNA-seq data have been deposited at the Gene Expression Omnibus^82^, under accession GSE289965. Raw and processed mass spectrometry proteomics data have been deposited at the ProteomeXchange Consortium via PRIDE^83^, under the identifier PDX060967.
- Time-lapse imaging videos and transmission electron microscopy images are available from the lead contact upon request.
- All original code has been deposited at Zenodo and is publicly available at 10.5281/zenodo.18393666 as of the date of publication.
- Any additional information required to reanalyze the data reported in this paper is available from the lead contact upon request.

## Acknowledgements

This work was supported by the Francis Crick Institute which receives its core funding from Cancer Research UK (CC2030, CC2052, CC2074), the UK Medical Research Council (CC2030, CC2052, CC2074), and the Wellcome Trust (CC2030, CC2052, CC2074). Work in the laboratory of KKN was supported by the Wellcome 221856/Z/20/Z and the Wellcome Human Developmental Biology Initiative 215116/Z/18/Z. The authors thank the members of the Crick Genomics Science Technology Platform (STP), the Crick Biological Research Facility and the Crick Genetic Modification Service. In particular, the authors would like to thank Lucy Collinson of the Crick Electron Microscopy STP, Sarah Maslen of the Crick Proteomics STP and Roger George and Chloë Roustan of the Crick Structural Biology STP. Additionally, we thank Agata Zielinska of the Crick Sex Chromosome Biology Laboratory for her advice and tutelage on handling oocytes and early embryos, and Daniel Snell of the Crick Genomics STP for his advice on sequencing. The authors would also like to thank Mads Lerdrup and Eva Hoffmann of the University of Copenhagen for their insightful discussions of the work. J. P. C. W was supported by a Crick-Imperial College London studentship funded by the Department of Chemistry at Imperial College London and the Francis Crick Institute.

## Author contributions

Conceptualization, L. J. W., J. M. A. T., M. T. B.;

Methodology, T. A., J. O., M. T. B., O. A. O.;

Formal Analysis, J. P. C. W., T. A.;

Investigation, J. P. C. W., T. A., A. M., A. E. W., D. L.;

Resources, O. A. O., J. M. A. T.;

Writing – Original Draft, J. P. C. W., L. J. W.;

Writing – Review & Editing, J. P. C. W., L. J. W., J. M. A. T., T. A., K. K. N.;

Visualisation, J. P. C. W.;

Supervision, L. J. W,. J. M. A. T., M. S., K. K. N.;

Project Administration, L. J. W., J. P. C. W., M. T. B.;

Funding Acquistion, L. J. W., J. M. A. T., K. K. N.;

## Methods

### Animal general husbandry

All animal procedures were performed in accordance with the relevant institutional and national guidelines and regulation, under the UK Home Office License Number P1234304. The animals were housed at the animal facility of The Francis Crick Institute, under a 12h light/12h dark cycle, in singly ventilated cages, at relative humidity between 45 and 65% and 20 – 24 °C. Reverse osmosis water and autoclaved pelleted food were available 24 h to the animals. Caging, bedding, nesting material and enrichments were autoclaved prior to use and changed routinely. The health of the animals was monitored by daily visual checks and weekly full health checks. Adult females and males were housed in separate cages in groups of 3-4 animals. Pups were genotyped for *Padi6^C663A^*and *Padi6* knock-out alleles using real time PCR with specific probes designed for each gene by Transnetyx (Cordova, TN). Only homozygous females between the ages of 8 and 11 weeks bearing the desired genetic mutations were used in experiments.

### Generation of Padi6 C663A mouse model

Three guide RNAs (sgRNAs) were designed using CRISPOR.org^84^ to cut in Exon 16 of the *Padi6* gene (Extended Data Figure 2, Supplementary Table 1). A 120bp ssODN was designed to be used with all three guides and containing the C663A (TGT>GCC) mutation, silent mutations for the respective guides, and 58-59 bp homology arms either side (Supplementary Table 2). C57BL/6J zygotes were electroporated using the Nepa21 square wave electroporator (Sonidel). 50 μL of the RNP complex together with the ssODN (1.2 μM/ 6 μM/ 8 μM was placed in the chamber of a CUY505P5 electrode, and the zygotes were electroporated using a poring pulse of 225 V (length: 1 ms, interval: 50 ms, #: 4, D. Rate: 10%, polarity: +) and a transfer pulse of 20 V (length: 50 ms, interval: 50 ms, #: 5, D. Rate: 40%, polarity: +/-). Real time PCR was performed by Transnetyx (Cordova, TN) on mouse ear-snips to identify eleven F0 mice preliminarily positive for the mutation. Amplicon PCR was performed on all founder mice spanning the entire ssODN and submitted for high-throughput sequencing using Illumina MiSeq sequencing 250 bp paired-end reads at the Crick Genomics Science Technology Platform (primers listed in Supplementary Table 3). Targeting with sgRNA-3 produced six F0 mice with perfect integration of the point mutations; three of these mice, with the highest percentage of the intended mutation, were crossed to C57BL/6J and produced two and six F1 mice in independent lines, identified as heterozygous for the mutation by Illumina MiSeq sequencing. Long read sequencing (Oxford Nanopore Technologies) was performed on 3.2 kb amplicons, generated by PCR, spanning the targeted insertion on a single F1 mouse from each line, no aberrant on-target mutations or deletions were found 1.5 kb upstream or downstream of the insertion in either case (primers listed in Supplementary Table 4). Copy number was evaluated using ddPCR on the same two F1 heterozygous mice and no evidence of random integrations of the transgene was found (primers listed in Supplementary Table 5).

### Generation of Padi6 knock-out mouse model

C57BL/6N *Padi6^tm1a(KOMP)Wtsl^*/MbpMmucd line (ID:48953) sperm was obtained from Mutant Mouse Resource & Research Centres (MMRC). The critical exon was deleted by inseminating oocytes from *PGK-Cre* mice by in vitro fertilization (IVF). After embryo transfer, *Padi6* mice with a lacZ tagged null allele (tm1b) were generated. Mice were genotyped using real time PCR with specific probes designed by Transnetyx (Cordova, TN).

### Generation of sterile XX^Y*^ mice

Sterile XX^Y*^ (X-attached-Y) mice were obtained by mating C57BL/6 females to XY* mice carrying a rearranged Y chromosome^85^. Recombination between an X and the Y* chromosome during male meiosis results in the X^Y*^ recombinant product, in which almost the entire Y* chromosome is attached to the X. Adult sterile XX^Y*^ (20 mg testis) mice are identified by testis palpation under short-term inhalation anaesthesia.

### Cloning and preparation of recombinant DNA

Primers were designed using SnapGene® software for In-Fusion cloning (primers listed in Supplementary Table 6). The coding sequences of mouse *Padi6* (Twist Bioscience), human *PADI6* (DNASU, #HsCD00297377), UHRF1 (Addgene, #142860^86^), CUL1 (donated by Dr Radoslav Enchev) and UBE2D2 (donated by Dr Katrin Rittinger) were amplified by PCR. The pcDNA3.1-Strep-Strep-TEV (donated by Dr Neil Q. McDonald^87^) and pcDNA3.1-3xFLAG (donated by Prof. Chris Schofield) and pcDNA3.1-HA (gift from Christopher A Walsh, Addgene 32530) vector backbones were linearized by PCR. The pcDNA3-MYC vector was linearised using EcoRI and NotI restriction enzymes. Amplified fragment inserts and linearised vectors were resolved by gel electrophoresis in a 1% Agarose TAE gel (100 V, 30 min) in TAE buffer and extracted using the PureLink™ Quick Gel Extraction Kit (Invitrogen, #K210012) following the manufacturer’s protocol. Fragment (∼50 ng) and linearised vector (∼50 ng) were combined with 2 µL 5X In-Fusion HD Enzyme (Takara Bio, #639650) and mixture made up to 10 µL with Nuclease Free Water. The mixture was incubated for 15 min at 50 °C and 5 µL transformed into 50 µL a *E. Coli* NEB® 5-alpha aliquot (New England Biolabs, #C2987H). Plasmids were sequenced by Sanger sequencing to confirm successful gene insertion and MaxiPrepped using the ZymoPURE II™ Plasmid Maxiprep kit (Zymo, #D4202) before transfection into Expi293™ cells or HEK-293T cells.

### Plasmid mutagenesis

Mutagenesis primers were designed to be ∼35 bp in length, with a melting temperature >78 °C with a GC content of ∼40% terminating in C or G (primers listed in Supplementary Table 7). Mutagenesis was performed by PCR following the Agilent QuickChange Protocol with PfuTurbo DNA polymerase (Agilent, #600250). After PCR amplification, 1 µL Dnp1 restriction enzyme (NEB, #R0176S) was added to the PCR mixture and the sample incubated at 37 °C for 1 h. After 1 h, 5 µL were transformed into a E. Coli NEB® 5-alpha aliquot (New England Biolabs, #C2987H). Plasmids were sequenced by Sanger sequencing to confirm presence of desired mutation and MaxiPrepped using the ZymoPURE II™ Plasmid Maxiprep kit (Zymo, #D4202) before transfection into Expi293™ or HEK-293T cells.

### Recombinant protein expression and purification

Expi293™ cells (Gibco™, #A14527) were incubated with 150 rpm agitation, 8% CO_2_, at 37 °C and diluted twice weekly. For transfection, cells were counted using Vi-CELL XR counter (Beckmann Coultier) and diluted to 2 x 10^6^ cells/mL in pre-warmed Expi293™ Expression Medium (ThermoFisher, #A1435101). 24 hours later, cells were transfected with MaxiPrepped recombinant DNA and PEI 25K™ (Polysciences, #23966-1) at a ratio of 1:3 (DNA: PEI) in Opti-MEM® (ThermoFisher, #11058021). Cultures were then incubated for 3 days before harvesting by centrifugation at 2000 rpm for 15 min at 4 °C and cell pellets flash frozen in liquid nitrogen and stored at -80 °C.

Frozen transfected Expi293™ cell pellets were re-suspended in purification buffer (50 mM Tris-HCl, pH 7.5, 200 mM NaCl, 1 mM TCEP) with cOmplete™ EDTA-free Protease Inhibitor Cocktail tablets (Merck, #73567200), 1 mg/mL Lysozyme (Sigma, #L6876-5G) and a micro-spatula of DNAse (Roche, #10104159001). Cells were lysed on ice by sonication and crude lysates centrifuged at 21,000 rpm for 45 min before being filtered through 0.45 µm PVDF filters. Cleared cell lysate was added to StrepTactin® XT 4Flow® resin (Iba, #2-5010-025) at a ratio of 2 mL packed beads per 500 mL Expi293 cell culture and incubated at 4 °C for 4 hours with gentle rotation. After 4 h, the unbound fraction was removed by gravity filtration and the resin was washed with purification buffer. Washed resin was re-suspended in twice its volume of purification buffer, TEV protease (produced in-house) added at a ratio of 2 mg protease per 500 mL Expi293 cell culture and incubated at 4 °C overnight with gentle rocking. The cleaved fraction was then removed by gravity filtration and the resin was again washed with purification buffer. Cleaved and wash fractions were combined and concentrated to 2 mL using a Vivaspin™ 20, 30000 MWCO ultrafiltration centrifugal concentrator (Sartorius, #10506282) and purified by size exclusion chromatography using a HiLoadTM 16/600 SuperdexTM 200 pg Gel Filtration Column (Cytivia, # 28989335), in purification buffer on an ÄKTApure™ system (Cytivia). Fractions were characterised by SDS-PAGE and pooled. Pooled fractions were concentrated again until no further decrease in volume was observed and stored at -80 °C.

### Intact-MS

Proteins were denatured and diluted to 1 µM at pH 2 with 2% (v/v) formic acid and 2% (v/v) acetonitrile. Denatured proteins were injected onto a C4 BEH 1.7 µm, 1 mm x 100 mm UPLC column (Waters™, #186005590) using an Acquity I class LC (Waters™). Proteins were eluted with a 15 min gradient of acetonitrile (2% v/v to 80% v/v) in 0.1% v/v formic acid using a flow rate of 50 µL/min. The analytical column outlet was directly interfaced via an electrospray ionisation source, with a time-of-flight BioAccord mass spectrometer (Waters™). Data was acquired over a *m/z* range of 300 – 8000, in positive ion mode with a cone voltage of 40 V. Scans were summed together manually and deconvoluted using MaxEnt1 (Masslynx, Waters™). Deconvoluted mass and intensity values were exported and plotted in GraphPad Prism.

### Nano differential scanning fluorimetry

Proteins were diluted to an approximate concentration of 0.2 mg/mL in purification buffer (50 mM Tris-HCl, pH 7.5, 200 mM NaCl, 1 mM TCEP). Melting experiments were performed using Prometheus™ NT.48 High Sensitivity Capillaries (NanoTemper, #PR-C006-200) on a Prometheus (NanoTemper) instrument with a 20 to 95 °C melt at 1 °C per minute. The 350/30 nm fluorescence ratio and first derivative of this ratio was plotted against temperature using GraphPad Prism. Melting temperatures were extracted as the first derivative peak in the melting curve calculated using the Panta Analysis software (NanoTemper). Experiment was conducted in triplicate.

### Mass Photometry

Proteins were diluted to an approximate concentration of 20 nM in purification buffer (50 mM Tris-HCl, pH 7.5, 200 mM NaCl, 1 mM TCEP). 2 µL of diluted protein was then added to 18 µL of PBS in the well of a gasket on a TwoMP instrument (Refeyn) and events recorded over the course of 1 min using the AquireMP software (Refeyn). Molecule sizes were calculated using the AnalyseMP software (Refeyn) using BSA (Thermo Scientific™, #23209), ADH (Sigma-Aldrich, #A7011) and Urease (Sigma-Aldrich, #94280) as standards. Mass normalised event histograms were produced using GraphPad Prism with bin sizes of 5 kDa.

### Mouse ovary RT-qPCR

Ovaries were homogenised using a 21G BD Microlance™ 3 needle (Camlab, #304432) and mRNA extracted using RNeasy® Mini Kit (Qiagen, #74104) following the manufacturer’s protocol. Extracted mRNA was reverse transcribed using the LunaScript® RT Super Mix Kit (NEB, #E3010S) following the manufacturer’s protocol. Transcribed mRNA was diluted in Nuclease Free Water to 5 ng/µL and RT-qPCR was performed using QuantiTect® qPCR primer assays (Qiagen) for *Padi6* (Mm_Padi6_1_SG, #QT00162750) and *B2m* (Mm_B2m_2_SG, #QT01149547) with the PowerUp™ SYBR™ Green Master Mix (Applied Biosystems™, #A25741) following the manufacturer’s protocol on a QuantStudio™ 12K Flex Real-Time PCR System (Applied Biosystems™) instrument following the standard PowerUp™ SYBR™ Green PCR program.

### Evaluation of female mouse fertlity

To evaluate fertility, females were mated over a period of at least 21 days with wild-type male stud mice. The number of pups born per litter was recorded. 14 wild-type females produced 14 litters, 8 *Padi6^C663A/C663A^*females produced 8 litters, and 5 *Padi6^-/-^* females did not produce any litters.

### Timed natural matings

To gather MII oocytes, females were time mated with sterile XX^Y*^ males, as previously described^85^. For zygotes and 2-cell embryos, females were time mated with wild-type C57BL/6J males. Natural matings were conducted as follows: females were placed into the cage of males at approximately 1700 h. The following morning, the presence of a vaginal plug was taken as evidence of successful mating and designated as Day 1 of gestation (E0.5).

### GV oocyte collection

Female mice between the ages of 8 and 10 weeks were culled, ovaries removed and placed in 2 mL of room temperature Medium2 (M2) (Sigma-Aldrich, #M7167) with 250 µM Dibutyryl cAMP (DBcAMP, Sigma-Aldrich, #D0260-5MG) in a 35 mm cell culture dish (Corning, #430165). Using a Leica MC80 dissecting microscope, ovarian follicles were repeatedly punctured with an 30G BD Microlance™ 3 needle (Camlab, #304432) to release oocytes. GV oocytes were identified by presence of a clear nucleus in the centre of a round cell. Oocytes were aspirated using a Stripper pipette with a 275 µm tip (Cooper Surgical, #MXL3-STR and #MXL3-275), and washed five times through droplets of M2 medium, followed by three washes in Dulbecco’s PBS (Gibco, #14190-094).

### MII oocyte collection

At approximately 1700 h on the day a vaginal plug was found, females were culled, and their uteri removed and placed in a 50 µL droplet of room temperature M2 medium (Sigma-Aldrich, #M7167) in a 60 mm dish (Corning, #430196). Using a Leica MC80 dissecting microscope, uteri were dissected using surgical forceps (Dumont, #0208-5-PO) and cumulus masses were aspirated using a Stripper pipette with a 275 µm tip (Cooper Surgical, #MXL3-STR and #MXL3-275). Cumulus cells were removed by incubation in 3 mg/mL Hyaluronidase (Sigma-Aldrich, #H4272-30G) in FHM for 5 seconds and washed five times in M2 medium, followed by three washes in Dulbecco’s PBS (Gibco, #14190-094).

### Collection of zygotes

At approximately 1700 h on the day a vaginal plug was found, females were culled, and oviducts were removed and placed in Embryo Max® FHM medium (Sigma Aldrich, #MR-024D) in a 60 mm dish (Corning, #430196). Under a Leica MC80 dissecting microscope, the infundibulum was torn using using surgical forceps (Dumont, #0208-5-PO) releasing the cumulus masses. Cumulus cells were removed by incubation in 3 mg/mL Hyaluronidase (Sigma-Aldrich, #H4272-30G) in FHM medium and zygotes were identified by the presence of one or two polar bodies, and clearly visible pronuclei. Zygotes were then washed five times in FHM medium, followed by three washes in Dulbecco’s PBS (Gibco, #14190-094).

### Collection of 2-cell embryos

Time mated females were culled at 0900 h on Day 2 of gestation and oviducts placed Embryo Max® FHM medium (Sigma Aldrich, #MR-024D) in a 60 mm dish (Corning, #430196). Using a Leica MC80 dissecting microscope, uteri were dissected using surgical forceps (Dumont, #0208-5-PO) and 2-cell embryos were aspirated using a Stripper pipette with a 275 µm tip (Cooper Surgical, #MXL3-STR and #MXL3-275). 2-cell embryos were then washed five times through droplets of FHM medium, followed by three times through droplets of Dulbecco’s PBS (Gibco, #14190-094).

### Dissociation of 2-cell embryos into blastomeres

For scRNA-seq, 2-cell embryos were dissociated into individual blastomeres. Briefly, after an initial wash in Embryo Max® FHM medium (Sigma Aldrich, #MR-024D), the zona was removed by brief incubation in 1X acidic Tryode’s solution (Sigma-Aldrich, #T1788) for approximately 1 min or until the zona pellucida had visibly disappeared. 2-cell embryos were then washed three times in FHM medium and incubated with 1X StemPro™ Accutase™ (Gibco™, #A1110501) for 5 min at room temperature. Following the incubation the embryos were washed in FHM medium and dissociated into single blastomeres by repeatedly aspirating and forcefully ejecting the embryo using a Stripper pipette with a 100 µm tip (Cooper Surgical, #MXL3-STR and #MXL3-100).

### Time-lapse imaging of zygote development to blastocyst

Mouse zygotes were cultured in pre-equilibrated Global medium (LifeGlobal, #LGGG-20) supplemented with 10% human serum albumin supplement (LifeGlobal, #GHSA-125) and overlaid with mineral oil (Origio, #ART-4008-5P). The embryos were cultured in EmbryoSlide+ culture dishes, using an EmbryoScope+ time-lapse incubator (VitroLife) at 37 °C, 5.5% carbon dioxide (CO_2_), over a period of five days. Time-lapse videos of embryo development were then analysed using Fiji. Pro-nuclear fading time was used to normalise the time at which the embryos reached the following stages: 2-cell, 4-cell, 8-cell, morula, cavity formation, and blastocyst. A total of 68 zygotes from 11 wild-type females, 84 mat-zygotes from 12 *Padi6^C663A/C663A^* females, 43 zygotes from 8 *Padi6^-/-^* females were imaged.

### Transmission electron microscopy

Oocytes were processed for electron microscopy using a Pelco BioWave Pro+ microwave (Ted Pella Inc, Redding, USA). Samples were fixed in in 2.5% (v/v) glutaraldehyde (Taab, #G002) / 4% (v/v) formaldehyde (Taab, #F017) in 0.1 M phosphate buffer (PB), pH 7.4, for one hour at RT. The steps including osmium, tannic acid, sodium sulphate and ethanol dehydration and 100% acetone up to and including 75% resin in acetone was performed in the Biowave, except for the phosphate buffer (PB) and dH_2_O wash steps, which consisted of two washes on the bench followed by two washes in the Biowave without vacuum at 250 W for 40 s each. Osmium and tannic acid steps were performed in the Biowave for 14 min under vacuum in 2 min cycles alternating with/without 100 W power. The SteadyTemp plate was kept at 21 °C. The sodium sulphate step was performed in the Biowave for 1min under vacuum with 100 W power with the SteadyTemo plate kept at 21 °C.

After primary fixation in 2.5% (v/v) glutaraldehyde (Taab, #G002) / 4% (v/v) formaldehyde (Taab, #F017) in 0.1 M phosphate buffer (pH 7.4), samples were stained with 1% (v/v) osmium tetroxide (Taab, #O012) / 1.5% (v/v) potassium ferricyanide (Sigma, #702587), washed in dH_2_O, incubated in 1% (w/v) tannic acid (Sigma, #702587) in 0.05 M PB, pH 7.4, washed in dH_2_O followed by 1% sodium sulphate in 0.05 M PB, pH 7.4 (Sigma). The samples were then washed in dH_2_O and dehydrated in a graded ethanol series (25%, 50%, 70%, 90%, and 100%, twice each), and in 100% acetone (3 times) at 250 W for 40 s without vacuum. Samples were gradually infiltrated with Epon 812 resin (Taab) with 25% 50% and 75% resin in acetone steps, at 250 W for 3 min each, with vacuum cycling (on/off at 30 sec intervals). Samples were transferred to 100% resin overnight before mounting in embedding moulds and polymerising at 60 °C for 48 h.

To confirm the integrity of the oocytes, the resin blocks were trimmed and scans were performed at either 50 kV / 4 W or 40 kV / 3 W, with no filter, 1601 projections and a pixel size of between 3.3 – 1.1 μm using an Xradia 510 Versa (Zeiss). The data was automatically reconstructed using Scout-and-Scan^TM^ Control System Reconstructor software (Zeiss) and viewed in TXM3DViewer software (Zeiss). With the integrity of the oocytes confirmed, 350 nm sections were cut with a diamond knife on an RMC Powertome Ultramicrotome (Boeckeler), stained with 1% toluidine blue in borax and viewed under a III RS light microscope (Carl Zeiss UK) to locate the correct region for ultra-thin sectioning for transmission electron microscopy. Seventy nm sections were collected onto 150 hexagonal copper grids, stained for 2 min with lead citrate and images obtained using a JEOL 1400 Flash transmission electron microscope with Matataki Flash sCMOS camera. Transmission electron microscopy images were not processed or adjusted, a total of 4 wild-type, 6 *Padi6^C663A/C663A^*, and 4 *Padi6^-/-^* GV oocytes were imaged.

### scRNA-sequencing and analysis

Three embryos from two independent females were processed for each genotype and stage, resulting in a total of six MII oocytes and six zygotes per genotype. The 2-cell embryos were dissociated into individual blastomeres and processed separately resulting in 12 single 2-cell blastomeres per genotype. MII oocytes, zygotes and 2-cell embryo blastomeres were dispensed into 12.5 µL CDS sorting solution from the Takara Smart-SeqHT kit (TakaraBio, #634439) and stored at -80 °C. Sequencing libraries were prepared using the Takara Smart-SeqHT Kit (Takara Bio, #634439) according to the manufacturer’s instructions. Briefly, cells were lysed and polyadenylated RNA captured in a total volume of 12.5 µL. One-Step first strand synthesis and double stranded cDNA amplification were carried out with 18 PCR cycles. Amplified cDNA was purified via a SPRISelect (Beckman Coulter, #B23317) 1X bead clean-up and eluted in 15 µL Elution Buffer. Nextera Dual Index libraries were prepared with the Nextera XT kit (Illumina, #FC-131-1096). Purified cDNA was normalised to 0.2 ng/µL input and tagmentation was carried out by adding 2.5 µL Tagment DNA Buffer + 1.25 µL Amplicon Tagment mix to 1.25 µL cDNA. The tagmentation reaction was run for 10 min at 55 °C. A final index PCR was performed after adding 2.5 µL Nextera compatible dual indices (at 10 µM) + 3.75 µL NPM PCR Mix.

Amplified libraries were purified via a SPRIselect (Beckman Coulter, #B23317) 0.9X bead clean-up. The quality and fragment size distributions of the purified libraries was assessed by a 4200 TapeStation Instrument (Agilent). Libraries were pooled and sequenced on the Illumina NovaSeq6000 in PE100 configuration to a read depth of at least 5M reads per sample.

Fastq files were mapped to the GRCm38 genome and converted to gene count files using Nextflow nf-core/rnaseq pipeline^88^ with parameters: aligner = star_rsem, genome = mm10. Transcriptomic data processing was performed using the Python package ScanPy^89^. Gene count files were imported, and the 24411 identified genes were filtered to those present in at least 6 cells with 3000 minimum counts resulting in a filtered dataset of 7551 genes. Counts were normalised to counts per million (CPM) and then converted to log_2_ values. PCA analysis was performed using ScanPy on the full dataset, as well as oocyte, zygote, and 2-cell split data. Principle components were extracted and plotted using GraphPad Prism. To identify dysregulated genes, for each stage, mean log_2_CPM ratios (test vs. wild-type) and q-values were calculated for each gene, extracted and plotted using GraphPad Prism. Those genes that had a log_2_CPM fold change < -0.5 or > 0.5 with a q-value < 0.05 were highlighted as down and upregulated respectively. GO analysis was performed using the Database for Annotation, Visualization and Integrated Discovery (DAVID)^90,91^, and Benjamini-Hochberg adjusted p-values extracted and plotted in GraphPad Prism. Over representation analysis comparing dysregulated genes with the transcriptomic database of mouse early embryo development (DBTMEE v2)^53^ was performed using Python, and p-values adjusted using the Benjamini-Hochberg procedure, log_2_(observed/expected) and -log_10_(corrected p-values) were extracted and plotted using GraphPad Prism. Transcript Z-scores were calculated using Python package ScanPy and plotted using the Python package matplotlib showing the mean of all genes belonging to each subset with the 95% confidence interval as a shaded area above and below the mean.

### Single oocyte and embryo proteomics

GV oocytes and 2-cell embryos were harvested from at least 2 different females per genotype. Samples were lysed in 5 µL of proteomics lysis buffer (100 mM TEAB, pH 8.5 (Sigma, # T7408-100 mL), 0.2% DDM, 10 ng/µl Trypsin (Waters, #90057)) with brief sonication (10 sec in standard water bath sonicator), and digested overnight at 37 °C. Following digestion samples were acidified and loaded onto Evotips (Evosep, #EV2013) for MS analyses using the standard preparation procedure apart from that sample loading was done by centrifugation at 400 g. Peptides were analysed using the Evosep+ 40SPD Whisper method (Evosep) on an Aurora Elite column (Ion Optics). Data were acquired on a timsTOF Ultra (Bruker) operated in DIA-PASEF mode with variable windows. Scan width was set between 100 and 1700 m/z, ion mobility window: 1/K0 0.64 – 1/KO 1.45, ramp time: 150 ms and accumulation time: 150 ms (data acquisition windows in Supplementary Table 8). Raw data were processed using Spectronaut 18 searching against the mouse Uniprot database (09/2022) using the default settings, with local normalisation and global imputation enabled.

Locally normalised and globally imputed proteomics data was analysed using Python. Differentially expressed proteins were identified as for the transcriptomic analysis. For each stage, log_2_ protein intensity ratios (test vs. wild-type) and q-values were calculated for each protein, extracted and plotted using GraphPad Prism. Those proteins that had a log_2_ protein intensity fold change < -0.5 or > 0.5 with a q-value < 0.05 were highlighted as down and upregulated respectively. The Search Tool for the Retrieval of Interacting Genes/Proteins (STRING) was used for gene ontology pathway enrichment analysis of proteomics data: https://string-db.org/^59^. Enriched terms were filtered to include only those with an enrichment strength > 0.5 or > 0.75, with false-discovery rate cutoffs as indicated in figure legends.

### Triton X-100 treatment of GV oocytes and analysis

GV oocytes from at least 2 females per genotype were characterised. GV oocytes were transferred into 500 µL of CPL-extraction buffer (20 mM HEPES, 100 mM KCl, 20 mM MgCl_2_, 3 mM EGTA, 0.1% Triton X-100), in a non-adherent Nunc cell culture dish (ThermoFisher, #176740) and incubated at room temperature for 10 min. After 10 min, oocytes were thoroughly washed through three wells of 500 μL of PBS followed by 8 x 50 µL droplets of PBS before being dispensed into 5 μL of proteomics lysis buffer and stored at -80 °C. Triton X-100 treated oocytes were then processed and analysed using the same single oocyte/embryo mass spectrometry workflow used for untreated samples.

Locally normalised and globally imputed proteomics data from Triton X-100 treated GV oocytes was analysed using Python as for intact GV oocytes. Proteins with a log_2_ protein intensity ratio (*Padi6^-/-^* vs. wild-type) < -0.5 and a q-value < 0.05 were classed as CPL-enriched. Over-representation analysis was performed as for the scRNA-sequencing data. STRING analysis was conducted as for proteomic data from intact GV oocytes and 2-cell embryos.

### Co-immunoprecipitation and western blotting

Human Embryonic kidney cells (HEK 293-T, Crick Cell Science Technology Platform, The Francis Crick Institute) were cultured in DMEM (Gibco™, #11965092) supplemented with 10% Foetal Bovine Serum (Sigma, #F7524-50ML) with Penicillin/Streptomycin (100 µg/mL, Gibco™, #15140122) in a 5% CO_2_ atmosphere at 37 °C. HEK 293T cell line was profiled by short tandem repeat analysis (STR) and tested negative for mycoplasma. 24 h prior to transfection, cells were seeded at 1 x 10^6^ cells per well of a Nunc™ cell-culture treated 6-well plate (ThermoScientific™, #140675). Transient transfections were done using Lipofectamine™ 2000 (Invitrogen™, #11668027) in 1 mL Opti-MEM per well (ThermoScientific™, #11058021) according to the manufacturer’s protocol. After 4 h transfection, Opti-MEM was replaced with pre-warmed DMEM (10% FBS, 100 µg/mL penicillin/streptomycin). At 48 h post-transfection, DMEM was discarded from the plate and cells were washed once with cold PBS buffer. PBS was aspirated, and cells lysed with immunoprecipitation buffer (50 mM HEPES, pH 8.0, 150 mM NaCl, 0.1% Triton X-100, 1mM DTT) on ice for 30 min. Lysates were clarified by centrifugation at 13,000 rpm for 15 min at 4 °C, and supernatants collected. Protein concentration was measured using Thermo Fisher BCA Protein assay (Thermo Fisher #23225). 20 μL of anti-FLAG® M2 magnetic bead (Millipore, #M8823) slurry was washed in immunoprecipitation buffer 3 times. For co-immunoprecipitation, 500 μg of transfected cell lysate in immunoprecipitation buffer was added to 20 μL of conjugated anti-FLAG antibody-bead slurry and incubated at 4 °C for 45 min. Beads were washed 4 times with 1 mL immunoprecipitation buffer and samples were eluted with 1X Laemli buffer (BioRad #1610747). Samples were then boiled and separated by polyacrylamide gel electrophoresis (SDS-PAGE) using mPAGE 4-20% Bis-Tris precast gels (Merck #MP4G15). Proteins were then transferred to a PVDF membrane (Cytiva Amersham Hybond0.45 µm, #10600023) via wet transfer. The membrane was blocked with 5% milk in phosphate-buffered saline with 0.1% Tween-20 (PBS-T). Membranes were then incubated with anti-FLAG-HRP (Merck #A8592) in PBS-T for 1h at room temperature (RT), anti-MYC (Invitrogen #SAB4301136) or HA (Cell Signalling #3724) in PBS-T overnight at 4 °C. Membranes were then washed three times with PBS-T and incubated with goat anti-rabbit HRP (Invitrogen #32460) in PBS-T for 1h at RT followed by three washes with PBS-T. Membranes were imaged with Pierce^TM^ ECL Plus (Thermo Fisher #32132) and imaged on a BioRad imager.

### AlphaFold prediction of interaction between PADI6 and UHRF1

Models of the interaction between dimeric human PADI6 (UniProt ID: Q6TGC4) and human UHRF1 (UniProt ID: Q96T88) were generated using AlphaFold 3^72,73^ via the AlphaFold Server (www.alphafoldserver.com). Full-length sequences for both proteins were used as input, with two copies of PADI6 and one copy of UHRF1. Predictions were performed using the default model parameters and five independent models were generated. The resulting models were visualized using UCSF ChimeraX (v1.11.1) and analysed alongside their predicted aligned error (PAE) heatmaps loaded in using the AlphaFold Error Plot tool. Predicted interactions were mapped onto the structure with the command *alphafold contacts* with a maximum distance of 4.0 Å. The predicted error of contacts are coloured from low error strong contacts to high error weak contacts, blue to yellow to red. Predicted aligned error (PAE) heatmaps were plotted using Python. For clarity in figures only one PADI6 molecule is shown.

### Quantification and Statistical Analysis

Bar charts were plotted using GraphPad Prism, with the black line representing the mean, and all data points shown. Unless otherwise stated, comparison of two groups was performed using GraphPad Prism using an unpaired two-tailed Student’s t-test, with Welch’s correction if variance was not equal between groups (ns: p > 0.05, *: p ≤ 0.05, **: p ≤ 0.01, ***: p ≤ 0.001, ****: p ≤ 0.0001). Comparison of Kaplan-Meier survival curves was performed using a log-rank Mantel-Cox test (ns: p > 0.05, *: p ≤ 0.05, **: p ≤ 0.01, ***: p ≤ 0.001, ****: p ≤ 0.0001). For scRNA-seq and single oocyte or embryo proteomics, a minimum of 6 total replicates harvested from at least 2 females were analysed for each condition. For volcano plots, p-values were calculated and corrected for multiple testing using the Benjamini-Hochberg method to produce q-values in Python; Jupyter notebooks for this analysis have been deposited to Zenodo (see Data and code Availability).

## Notes

### Competing Interest Statement

The authors have declared no competing interest.

### Summary of Updates

Figure 6 revised to include pairwise coIP data Changes from use of p values to q values for analysis of scRNA-seq and proteomics data sets

## References

1. Li, L., Zheng, P. & Dean, J. Maternal control of early mouse development. Development 137, 859–870 (2010).

2. Zhang, K. & Smith, G. W. Maternal control of early embryogenesis in mammals. Reprod. Fertil. Dev. 27, 880 (2015).

3. Wu, D. & Dean, J. Maternal factors regulating preimplantation development in mice. Curr. Top. Dev. Biol. 140, 317–340 (2020).

4. Tadros, W. & Lipshitz, H. D. The maternal-to-zygotic transition: a play in two acts. Development 136, 3033–3042 (2009).

5. Jukam, D., Shariati, S. A. M. & Skotheim, J. M. Zygotic Genome Activation in Vertebrates. Dev. Cell 42, 316–332 (2017).

6. Schulz, K. N. & Harrison, M. M. Mechanisms regulating zygotic genome activation. Nat. Rev. Genet. 20, 221–234 (2019).

7. Yurttas, P. et al. Role for PADI6 and the cytoplasmic lattices in ribosomal storage in oocytes and translational control in the early mouse embryo. Development 135, 2627–2636 (2008).

8. Kan, R. et al. Regulation of mouse oocyte microtubule and organelle dynamics by PADI6 and the cytoplasmic lattices. Dev. Biol. 350, 311–322 (2011).

9. Yu, X. J. et al. The subcortical maternal complex controls symmetric division of mouse zygotes by regulating F-actin dynamics. Nat. Commun. 5, 4887 (2014).

10. Gao, Z. et al. Zbed3 participates in the subcortical maternal complex and regulates the distribution of organelles. J. Mol. Cell Biol. 10, 74–88 (2018).

11. Rezaei, M. et al. Novel pathogenic variants in NLRP7, NLRP5, and PADI6 in patients with recurrent hydatidiform moles and reproductive failure. Clin. Genet. 99, 823–828 (2021).

12. Dong, J. et al. Novel biallelic mutations in PADI6 in patients with early embryonic arrest. J. Hum. Genet. 1–9 (2022) doi:10.1038/s10038-021-00998-8.

13. Xu, Y. et al. Novel Homozygous PADI6 Variants in Infertile Females with Early Embryonic Arrest. Front. Cell Dev. Biol. 10, 1–8 (2022).

14. Xu, Y. et al. Mutations in PADI6 Cause Female Infertility Characterized by Early Embryonic Arrest. Am. J. Hum. Genet. 99, 744–752 (2016).

15. Zheng, W. et al. New biallelic mutations in PADI6 cause recurrent preimplantation embryonic arrest characterized by direct cleavage. J. Assist. Reprod. Genet. 37, 205–212 (2020).

16. Liu, J. et al. Two novel mutations in PADI6 and TLE6 genes cause female infertility due to arrest in embryonic development. J. Assist. Reprod. Genet. 38, 1551–1559 (2021).

17. Wang, X. et al. Novel mutations in genes encoding subcortical maternal complex proteins may cause human embryonic developmental arrest. Reprod. Biomed. Online 36, 698–704 (2018).

18. Maddirevula, S. et al. The human knockout phenotype of PADI6 is female sterility caused by cleavage failure of their fertilized eggs. Clin. Genet. 91, 344–345 (2017).

19. Zhang, T., Liu, P., Yao, G., Zhang, X. & Cao, C. A complex heterozygous mutation in PADI6 causes early embryo arrest: A case report. Front. Genet. 13, 1104085 (2023).

20. Qian, J. et al. Biallelic PADI6 variants linking infertility, miscarriages, and hydatidiform moles. Eur. J. Hum. Genet. 26, 1007–1013 (2018).

21. Eggermann, T. et al. Trans-acting genetic variants causing multilocus imprinting disturbance (MLID): common mechanisms and consequences. Clin. Epigenetics 14, 1–17 (2022).

22. Eggermann, T., Kadgien, G., Begemann, M. & Elbracht, M. Biallelic PADI6 variants cause multilocus imprinting disturbances and miscarriages in the same family. Eur. J. Hum. Genet. 29, 575–580 (2021).

23. Begemann, M. et al. Maternal variants in NLRP and other maternal effect proteins are associated with multilocus imprinting disturbance in offspring. J. Med. Genet. 55, 497–504 (2018).

24. Cubellis, M. V. et al. Loss-of-function maternal-effect mutations of PADI6 are associated with familial and sporadic Beckwith-Wiedemann syndrome with multi-locus imprinting disturbance. Clin. Epigenetics 12, 139 (2020).

25. Tannorella, P. et al. Germline variants in genes of the subcortical maternal complex and Multilocus Imprinting Disturbance are associated with miscarriage / infertility or Beckwith – Wiedemann progeny. Clin. Epigenetics 14, 1–7 (2022).

26. Demond, H. et al. A KHDC3L mutation resulting in recurrent hydatidiform mole causes genome-wide DNA methylation loss in oocytes and persistent imprinting defects post-fertilisation. Genome Med. 11, 1–14 (2019).

27. Capco, D. G., Gallicano, G. I., McGaughey, R. W., Downing, K. H. & Larabell, C. A. Cytoskeletal sheets of mammalian eggs and embryos: A lattice-like network of intermediate filaments. Cell Motil. Cytoskeleton 24, 85–99 (1993).

28. McGaughey, R. W. & Capco, D. G. Specialized cytoskeletal elements in mammalian eggs: Structural and biochemical evidence for their composition. Cell Motil. Cytoskeleton 13, 104–111 (1989).

29. Ian Gallicano, G., Larabell, C. A., McGaughey, R. W. & Capco, D. G. Novel cytoskeletal elements in mammalian eggs are composed of a unique arrangement of intermediate filaments. Mech. Dev. 45, 211–226 (1994).

30. Jentoft, I. M. A. et al. Mammalian oocytes store proteins for the early embryo on cytoplasmic lattices. Cell 186, 5308–5327.e25 (2023).

31. Cheng, S. et al. Mammalian oocytes store mRNAs in a mitochondria-associated membraneless compartment. 262, (2022).

32. Zaffagnini, G. et al. Mouse oocytes sequester aggregated proteins in degradative super-organelles. Cell 187, 1109–1126.e21 (2024).

33. Liu, X. et al. Role for PADI6 in securing the mRNA-MSY2 complex to the oocyte cytoplasmic lattices. Cell Cycle 16, 360–366 (2017).

34. Williams, J. P. C. & Walport, L. J. PADI6: What we know about the elusive fifth member of the peptidyl arginine deiminase family. Philos. Trans. R. Soc. B Biol. Sci. 378, 20220242 (2023).

35. Gao, Y. et al. Protein Expression Landscape of Mouse Embryos during Pre-implantation Development. Cell Rep. 21, 3957–3969 (2017).

36. Wright, P. W. et al. ePAD, an oocyte and early embryo-abundant peptidylarginine deiminase-like protein that localizes to egg cytoplasmic sheets. Dev. Biol. 256, 74–89 (2003).

37. Esposito, G. et al. Peptidylarginine deiminase (PAD) 6 is essential for oocyte cytoskeletal sheet formation and female fertility. Mol. Cell. Endocrinol. 273, 25–31 (2007).

38. Horibata, S., Coonrod, S. A. & Cherrington, B. D. Role of PADs in Disease and Female Reproduction. J. Reprod. Dev. 58, (2012).

39. Witalisom, E., Thompson, R. & Hofseth, L. Protein Arginine Deiminases and Associated Cirtullination. Curr Drug Targets 16, 700–710 (2015).

40. Lewallen, D. M. et al. Chemical Proteomic Platform to Identify Citrullinated Proteins. ACS Chem. Biol. 10, 2520–2528 (2015).

41. Mondal, S. & Thompson, P. R. Protein Arginine Deiminases (PADs): Biochemistry and Chemical Biology of Protein Citrullination. Acc. Chem. Res. 52, 818–832 (2019).

42. Rebak, A. S. et al. A quantitative and site-specific atlas of the citrullinome reveals widespread existence of citrullination and insights into PADI4 substrates. Nat. Struct. Mol. Biol. 31, 977–995 (2024).

43. Taki, H. et al. Purification of enzymatically inactive peptidylarginine deiminase type 6 from mouse ovary that reveals hexameric structure different from other dimeric isoforms. Adv. Biosci. Biotechnol. 02, 304–310 (2011).

44. Ranaivoson, F. M. et al. Crystal structure of human peptidylarginine deiminase type VI (PAD6) provides insights into its inactivity. IUCrJ 11, 395–404 (2024).

45. Williams, J. P. C. et al. Structural insight into the function of human peptidyl arginine deiminase 6. Comput. Struct. Biotechnol. J. 23, 3258–3269 (2024).

46. Arita, K. et al. Structural basis for Ca2+-induced activation of human PAD4. Nat. Struct. Mol. Biol. 11, 777–783 (2004).

47. Slade, D. J. et al. Protein arginine deiminase 2 binds calcium in an ordered fashion: Implications for inhibitor design. ACS Chem. Biol. 10, 1043–1053 (2015).

48. Ranaivoson, F. M. et al. Crystal structure of human peptidylarginine deiminase type VI (PAD6) provides insights into its inactivity. IUCrJ 11, 395–404 (2024).

49. Rechiche, O., Lee, T. V. & Lott, J. S. Structural characterization of human peptidyl-arginine deiminase type III by X-ray crystallography. Acta Crystallogr. Sect. F Struct. Biol. Commun. 77, 334–340 (2021).

50. Douglas, C. et al. CRISPR-Cas9 effectors facilitate generation of single-sex litters and sex-specific phenotypes. Nat. Commun. 12, 6926 (2021).

51. Vonnahme, K. A., Wilson, M. E., Foxcroft, G. R. & Ford, S. P. Impacts on conceptus survival in a commercial swine herd. J. Anim. Sci. 80, 553–559 (2002).

52. Wimsatt, W. A. Some Comparative Aspects of Implantation. Biol. Reprod. 12, 1–40 (1975).

53. Park, S.-J., Shirahige, K., Ohsugi, M. & Nakai, K. DBTMEE: a database of transcriptome in mouse early embryos. Nucleic Acids Res. 43, D771–D776 (2015).

54. Giaccari, C. et al. A maternal-effect *Padi6* variant causes nuclear and cytoplasmic abnormalities in oocytes, as well as failure of epigenetic reprogramming and zygotic genome activation in embryos. Genes Dev. 38, 131–150 (2024).

55. Giaccari, C., Cecere, F., Argenziano, L., Pagano, A. & Riccio, A. New insights into oocyte cytoplasmic lattice-associated proteins. Trends Genet. 40, 880–890 (2024).

56. Yang, G., Xin, Q. & Dean, J. Degradation and translation of maternal mRNA for embryogenesis. Trends Genet. 40, 238–249 (2024).

57. Ye, Z. et al. One-Tip enables comprehensive proteome coverage in minimal cells and single zygotes. Nat. Commun. 15, 2474 (2024).

58. Israel, S. et al. An integrated genome-wide multi-omics analysis of gene expression dynamics in the preimplantation mouse embryo. Sci. Rep. 9, 13356 (2019).

59. Szklarczyk, D. et al. The STRING database in 2021: customizable protein–protein networks, and functional characterization of user-uploaded gene/measurement sets. Nucleic Acids Res. 49, D605–D612 (2021).

60. Tashiro, F. et al. Maternal-effect gene Ces5/Ooep/Moep19/Floped is essential for oocyte cytoplasmic lattice formation and embryonic development at the maternal-zygotic stage transition. Genes Cells 15, 813–828 (2010).

61. Kim, B., Kan, R., Anguish, L., Nelson, L. M. & Coonrod, S. A. Potential role for MATER in cytoplasmic lattice formation in murine oocytes. PLoS ONE 5, 1–8 (2010).

62. Qin, D. et al. The subcortical maternal complex protein Nlrp4f is involved in cytoplasmic lattice formation and organelle distribution. Development 146, dev.183616 (2019).

63. Uemura, S. et al. UHRF1 is essential for proper cytoplasmic architecture and function of mouse oocytes and derived embryos. Life Sci. Alliance 6, e202301904 (2023).

64. Li, L., Baibakov, B. & Dean, J. A Subcortical Maternal Complex Essential for Preimplantation Mouse Embryogenesis. Dev. Cell 15, 416–425 (2008).

65. Mahadevan, S. et al. Maternally expressed NLRP2 links the subcortical maternal complex (SCMC) to fertility, embryogenesis and epigenetic reprogramming. Sci. Rep. 7, (2017).

66. Monk, D., Sanchez-Delgado, M. & Fisher, R. NLRPS, the subcortical maternal complex and genomic imprinting. Reproduction 154, R161–R170 (2017).

67. Han, J. et al. NLRP7 participates in the human subcortical maternal complex and its variants cause female infertility characterized by early embryo arrest. J. Mol. Med. 101, 717–729 (2023).

68. Li, Y. & Hao, B. Structural Basis of Dimerization-dependent Ubiquitination by the SCFFbx4 Ubiquitin Ligase. J. Biol. Chem. 285, 13896–13906 (2010).

69. Zheng, N. et al. Structure of the Cul1–Rbx1–Skp1–F boxSkp2 SCF ubiquitin ligase complex. Nature 416, 703–709 (2002).

70. Cardozo, T. & Pagano, M. The SCF ubiquitin ligase: insights into a molecular machine. Nat. Rev. Mol. Cell Biol. 5, 739–751 (2004).

71. Petroski, M. D. & Deshaies, R. J. Mechanism of Lysine 48-Linked Ubiquitin-Chain Synthesis by the Cullin-RING Ubiquitin-Ligase Complex SCF-Cdc34. Cell 123, 1107–1120 (2005).

72. Abramson, J. et al. Accurate structure prediction of biomolecular interactions with AlphaFold 3. Nature 630, 493–500 (2024).

73. Jumper, J. et al. Highly accurate protein structure prediction with AlphaFold. Nature 596, 583–589 (2021).

74. Goddard, T. D. et al. UCSF ChimeraX: Meeting modern challenges in visualization and analysis. Protein Sci. 27, 14–25 (2018).

75. Meng, E. C. et al. UCSF CHIMERAX : Tools for structure building and analysis. Protein Sci. 32, e4792 (2023).

76. Sies, H., Berndt, C. & Jones, D. P. Oxidative Stress. Annu. Rev. Biochem. 86, 715–748 (2017).

77. Qian, J. et al. Biallelic PADI6 variants linking infertility, miscarriages, and hydatidiform moles. Eur. J. Hum. Genet. 26, 1007–1013 (2018).

78. Devriendt, K. Hydatidiform mole and triploidy: the role of genomic imprinting in placental development. Hum. Reprod. Update 11, 137–142 (2005).

79. Xu, Y. et al. Mutations in PADI6 Cause Female Infertility Characterized by Early Embryonic Arrest. Am. J. Hum. Genet. 99, 744–752 (2016).

80. Neeli, I., Khan, S. N. & Radic, M. Histone Deimination As a Response to Inflammatory Stimuli in Neutrophils. J. Immunol. 180, 1895–1902 (2008).

81. Christophorou, M. A. et al. Citrullination regulates pluripotency and histone H1 binding to chromatin. Nature 507, 104–108 (2014).

82. Edgar, R. Gene Expression Omnibus: NCBI gene expression and hybridization array data repository. Nucleic Acids Res. 30, 207–210 (2002).

83. Perez-Riverol, Y. et al. The PRIDE database resources in 2022: a hub for mass spectrometry-based proteomics evidences. Nucleic Acids Res. 50, D543–D552 (2022).

84. Concordet, J.-P. & Haeussler, M. CRISPOR: intuitive guide selection for CRISPR/Cas9 genome editing experiments and screens. Nucleic Acids Res. 46, W242–W245 (2018).

85. Burgoyne, P. S., Mahadevaiah, S. K., Perry, J., Palmer, S. J. & Ashworth, A. The Y* rearrangement in mice: new insights into a perplexing PAR. Cytogenet. Genome Res. 80, 37–40 (1998).

86. Joung, J. et al. A transcription factor atlas of directed differentiation. Cell 186, 209–229.e26 (2023).

87. Earl, C. P. et al. Capture, mutual inhibition and release mechanism for aPKC–Par6 and its multisite polarity substrate Lgl. Nat. Struct. Mol. Biol. 10.1038/s41594-024-01425-0 (2025) doi:10.1038/s41594-024-01425-0.

88. Harshil Patel et al. nf-core/rnaseq: nf-core/rnaseq v3.18.0 - Lithium Lynx. Zenodo 10.5281/ZENODO.1400710 (2024).

89. Wolf, F. A., Angerer, P. & Theis, F. J. SCANPY: large-scale single-cell gene expression data analysis. Genome Biol. 19, 15 (2018).

90. Huang, D. W., Sherman, B. T. & Lempicki, R. A. Systematic and integrative analysis of large gene lists using DAVID bioinformatics resources. Nat. Protoc. 4, 44–57 (2009).

91. Sherman, B. T. et al. DAVID: a web server for functional enrichment analysis and functional annotation of gene lists (2021 update). Nucleic Acids Res. 50, W216–W221 (2022).

92. Sievers, F. et al. Fast, scalable generation of high-quality protein multiple sequence alignments using Clustal Omega. Mol. Syst. Biol. 7, 539 (2011).

93. Rodríguez-Nuevo, A. et al. Oocytes maintain ROS-free mitochondrial metabolism by suppressing complex I. Nature 607, 756–761 (2022).

