## Supplementary Information for "Dissecting PADI6 function defines oocyte cytoplasmic lattices as regulatory hubs for fundamental cellular processes"

**Figure S1**. Characterisation of mPADI6$^{C663A}$ and generation of transgenic mouse lines. Related to Figure 1.

**Figure S2.** *Padi6* mouse models. Related to Figure 1.

**Figure S3**. CPLs are present in similar abundance and morphology in *Padi6*$^{C663A/C663A}$ females compared to wild-type. Related to Figure 2.

**Figure S4.** Transcriptomic analysis of MII oocytes, zygotes and 2-cell embryos from wild-type, *Padi6*$^{C663A/C663A}$ and *Padi6*$^{-/-}$ females. Related to Figure 3.

**Figure S5.** Proteomic analysis of GV oocytes and 2-cell embryos from wild-type, *Padi6*$^{C663A/C663A}$ and *Padi6*$^{-/-}$ females. Related to Figure 4.

**Figure S6.** Analysis of CPL-enriched proteins. Related to Figures 5 and 6.

**Figure S7.** AlphaFold Server predicted interaction between human UHRF1 and human PADI6. Related to Figure 6.

**Table S1.** Padi6 C663A generation guide RNA sequences

**Table S2.** Padi6 C663A ssODN sequence

**Table S3.** Primer Sequences for Padi6 C663A genotyping by MiSeq

**Table S4.** Primer Sequences for Padi6 C663A long read sequencing

**Table S5.** Primer Sequences for Padi6 C663A ddPCR

**Table S6.** In-Fusion cloning primers

**Table S7.** Mutagenesis primers

**Table S8.** Single oocyte/embryo DIA PASEF data acquisition windows

**Figure S8.** Uncut western blot images with replicates for Figure 6C

**Figure S9.** Uncut western blot images with replicates for Figure 6G

**Figure S10.** Uncut western blot images with replicates for Figure 6I

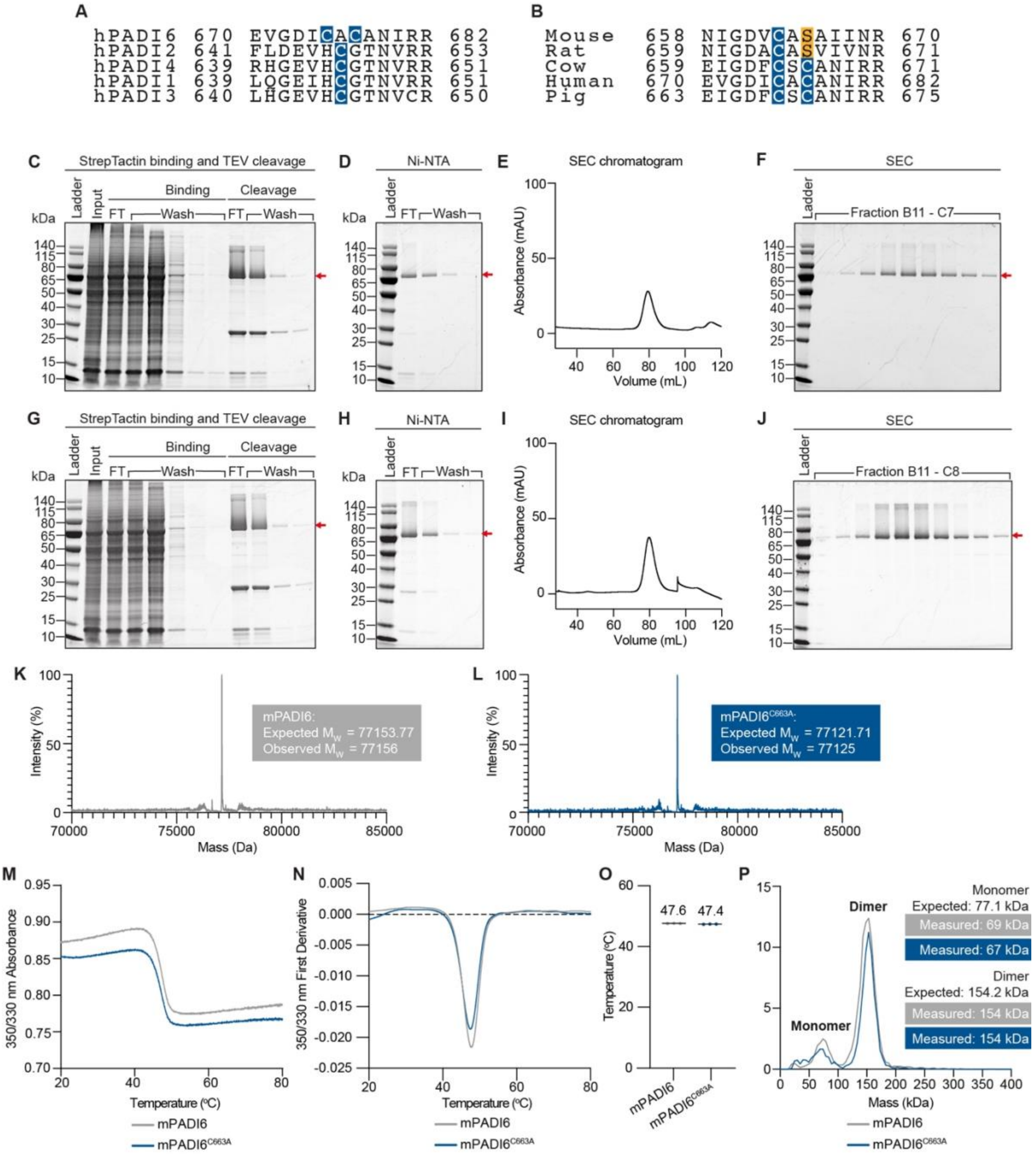


**Figure S1. Characterisation of mPADI6$^{C663A}$ and generation of transgenic mouse lines. Related to Figure 1.**

(A) Sequence alignments of human PADI key catalytic cysteine residues. The key catalytic cysteines conserved in human PADIs 1 to 4 are highlighted. The two possible catalytic cysteines in PADI6 are highlighted. Sequence alignment produced using Clustal Omega[92].

(B) Sequence alignments of the mouse, rat, cow, human, and pig PADI6 active site. Only the first of the cysteines is conserved in multiple species, C663 in mouse PADI6. Sequence alignment produced using Clustal Omega[92].

(C) SDS-PAGE gel after isolation of Strep-Strep-TEV-mPADI6 from transiently transfected Expi293 cells, and subsequent on-resin cleavage of mPADI6 with His-TEV protease. FT = flow through.

(D) SDS-PAGE gel of mPADI6 sample before and after application to Ni-NTA resin to remove His-TEV protease.

(E) Chromatogram of size-exclusion chromatography (SEC) purification of mPADI6.

(F) SDS-PAGE gel of mPADI6 SEC purification fractions. All characterised fractions combined and taken forward.

(G) As for (A) with mPADI6$^{C663A}$.

(H) As for (B) with mPADI6$^{C663A}$.

(I) As for (C) with mPADI6$^{C663A}$.

(J) As for (D) with mPADI6$^{C663A}$.

(K) Intact-MS spectrum of mPADI6.

(L) Intact-MS spectrum of mPADI6$^{C663A}$.

(M) NanoDSF melting spectra of mPADI6 and mPADI6$^{C663A}$. Absorbance was measured at 350 and 330 nm and the ratio plotted. Experiment performed in triplicate; line represents mean of 3 replicates.

(N) First derivative of 350 nm by 330 nm absorbance ratio shown in (K). Experiment performed in triplicate; line represents mean of 3 replicates.

(O) NanoDSF determined melting temperature of mPADI6 and mPADI6$^{C663A}$. Experiment performed in triplicate.

(P) Mass photometry histograms of mPADI6 and mPADI6$^{C663A}$ showing that both proteins predominantly exist as dimers in solution in vitro. Histogram bin size = 5 kDa.

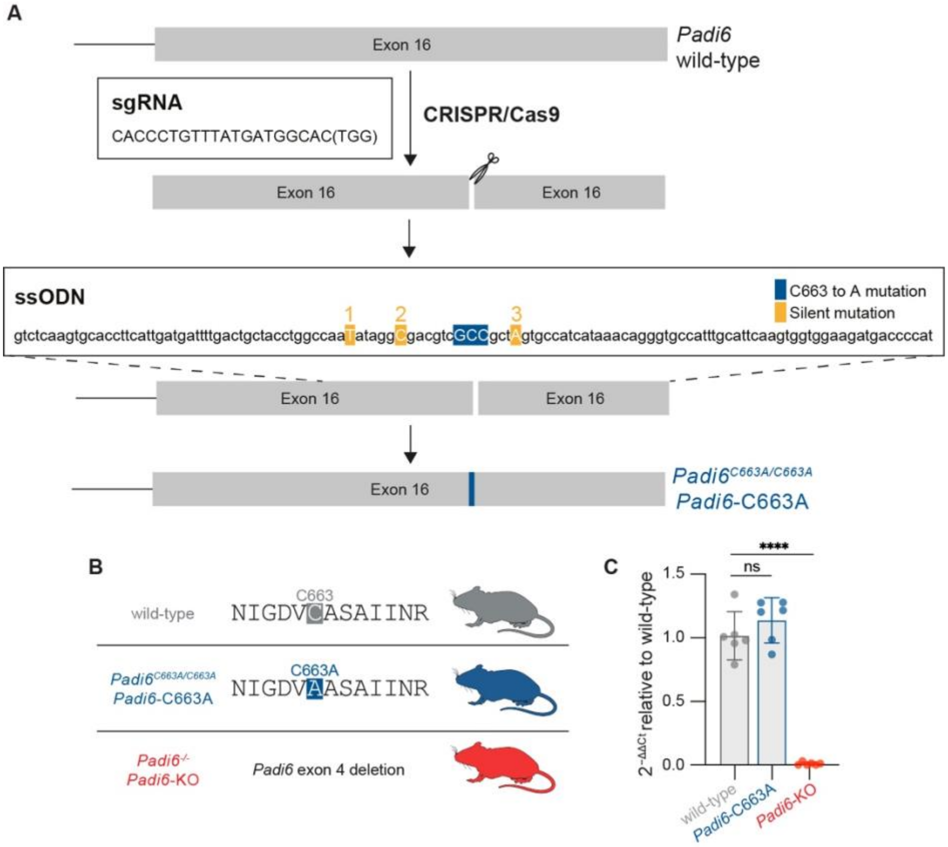


**Figure S2. *Padi6* mouse models. Related to Figure 1**

(A) Schematic representation of *Padi6*$^{C663A}$ mouse line generation. The successful sgRNA-3 is shown, which was electroporated with the RNP complex and ssODN into zygotes. After the CRISPR/Cas9 mediated double-strand DNA cleavage within exon 16 of the *Padi6* gene, DNA was repaired using a single ssODN mutating the wild-type TGT codon to GCC (C663 to A). Three silent mutations were included in the ssODN to mutate the PAM sequence of each of the guides. Silent mutation 3 mutates the PAM of sgRNA-3 to prevent re-cutting.

(B) Cartoon schematic depicting the *Padi6*$^{C663A/C663A}$ and *Padi6*$^{-/-}$ mouse lines.

(C) Relative *Padi6* expression levels in ovaries from 8-week-old wild-type, *Padi6*$^{C663A/C663A}$ (*Padi6*-C663A) and *Padi6*$^{-/-}$ (*Padi6*-KO) females determined by qRT-PCR. The data were normalised to *B2m* and the relative expression of *Padi6* determined compared to wild-type ovaries. Experiment was performed in technical triplicate, on ovaries from two different females of each genetic background. All six replicate values were plotted for each genetic background and the mean shown by a black line. Unpaired two-tailed Student's t-test; *$P \leq 0.05$, **$P \leq 0.01$, ***$P \leq 0.001$, ****$P \leq 0.0001$, ns ($P > 0.05$).

A

| | GV oocytes imaged | CPLs present |
|---|---|---|
| wild-type | 4 | 3/4 |
| *Padi6*-C663A | 6 | 5/6 |
| *Padi6*-KO | 4 | 0/4 |

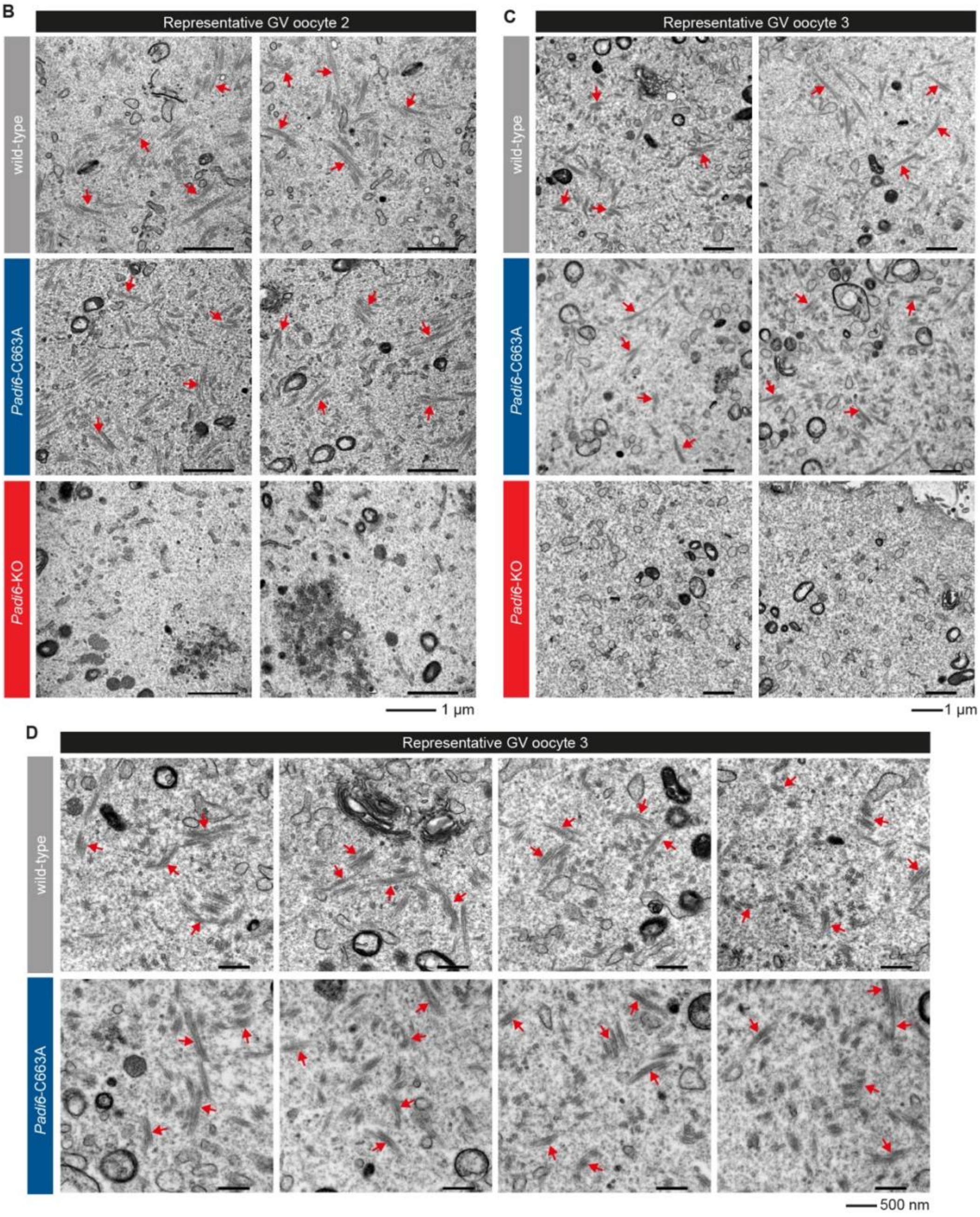


**Figure S3. CPLs are present in similar abundance and morphology in *Padi6*$^{C663A/C663A}$ females compared to wild-type. Related to Figure 2.**

(A) Summary of TEM imaging of germinal vesicle oocytes (GV) from wild-type, *Padi6*$^{C663A/C663A}$ (*Padi6*-C663A) and *Padi6*$^{-/-}$ (*Padi6*-KO) females.

(B and C) TEM images of representative GV oocytes 2 and 3 from wild-type, *Padi6*-C663A and *Padi6*-KO females showing the CPLs are present in oocytes from wild-type and *Padi6*-C663A females, but not *Padi6*-KO females. Representative CPLs are indicated by red arrows. Scale bar = 1 µm.

(D) Higher magnification TEM images of wild-type and *Padi6*-C663A representative GV oocyte 3 showing CPLs are similar morphology in wild-type and *Padi6*-C663A GV oocytes. Representative CPLs are indicated by red arrows. Scale bar = 500 nm.

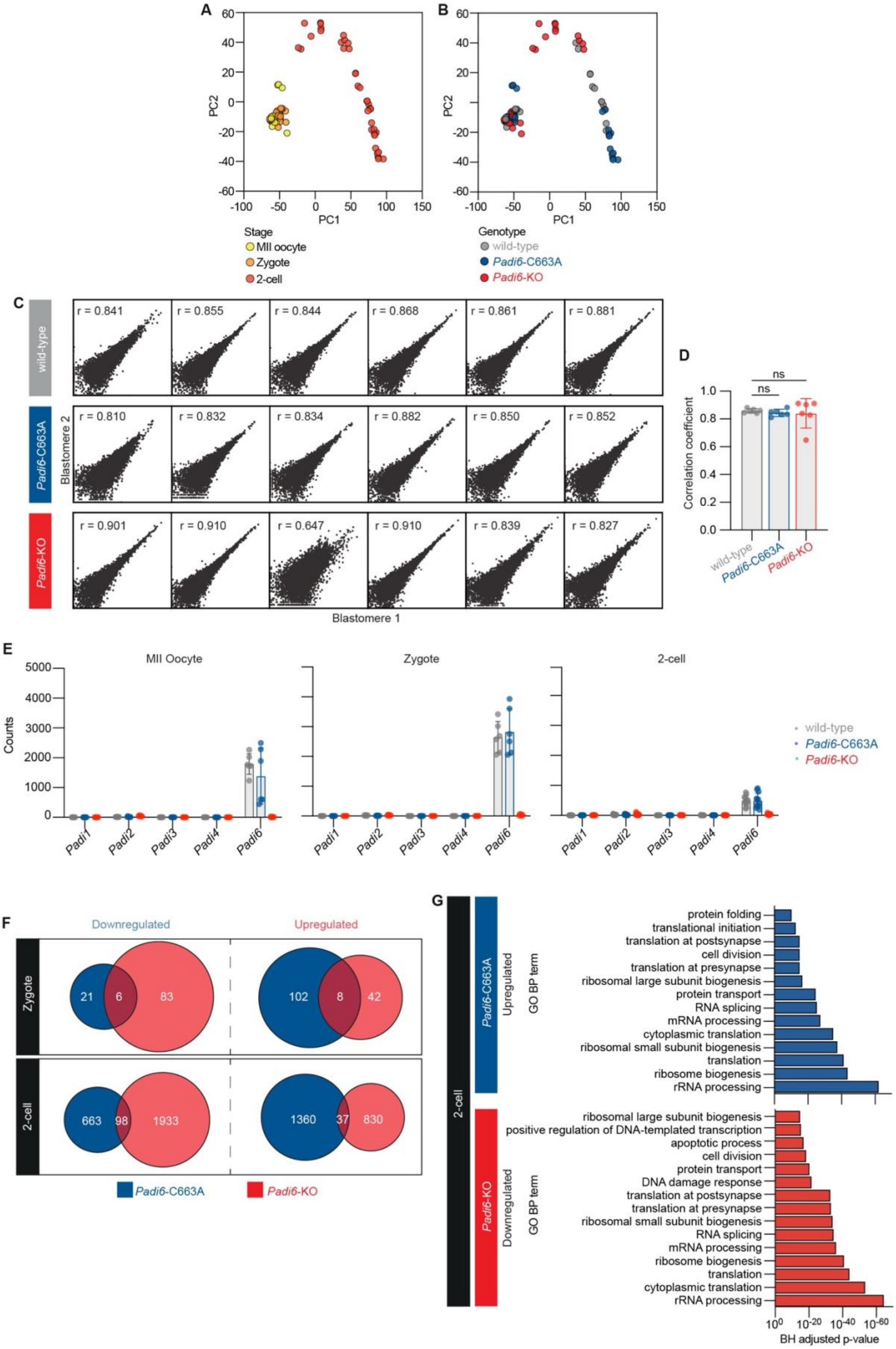
A
B
PC1
PC2
Stage
MII oocyte
Zygote
2-cell
Genotype
wild-type
Padi6-C663A
Padi6-KO
C
wild-type
Padi6-C663A
Padi6-KO
r = 0.841
r = 0.855
r = 0.844
r = 0.868
r = 0.861
r = 0.881
r = 0.810
r = 0.832
r = 0.834
r = 0.882
r = 0.850
r = 0.852
r = 0.901
r = 0.910
r = 0.647
r = 0.910
r = 0.839
r = 0.827
Blastomere 1
Blastomere 2
D
Correlation coefficient
ns
ns
E
MII Oocyte
Zygote
2-cell
Counts
Padi1
Padi2
Padi3
Padi4
Padi6
F
Downregulated
Upregulated
Zygote
2-cell
21
6
83
102
8
42
663
98
1933
1360
37
830
Padi6-C663A
Padi6-KO
G
2-cell
Padi6-C663A
Upregulated
GO BP term
protein folding
translational initiation
translation at postsynapse
cell division
translation at presynapse
ribosomal large subunit biogenesis
protein transport
RNA splicing
mRNA processing
cytoplasmic translation
ribosomal small subunit biogenesis
translation
ribosome biogenesis
rRNA processing
Padi6-KO
Downregulated
GO BP term
ribosomal large subunit biogenesis
positive regulation of DNA-templated transcription
apoptotic process
cell division
protein transport
DNA damage response
translation at postsynapse
translation at presynapse
ribosomal small subunit biogenesis
RNA splicing
mRNA processing
ribosome biogenesis
translation
cytoplasmic translation
rRNA processing
BH adjusted p-value

**Figure S4. Transcriptomic analysis of MII oocytes, zygotes and 2-cell embryos from wild-type, *Padi6*$^{C663A/C663A}$ and *Padi6*$^{-/-}$ females. Related to Figure 3.**

(A) PCA plot for all cells in the scRNA-seq dataset, coloured by developmental stage.

(B) PCA plot for all cells in the scRNA-seq dataset, coloured by genotype. *Padi6*-C663A = oocyte or embryo from *Padi6*$^{C663A/C663A}$ females, *Padi6*-KO = oocyte or embryo from *Padi6*$^{-/-}$ females.

(C) We did not observe significant differences between transcript levels in individual blastomeres from the same 2-cell embryo. Scatter plot of $log_2$ CPM values for sister blastomeres of 2-cell embryos from wild-type, *Padi6*-C663A and *Padi6*-KO females. Pearson correlation coefficient for each 2-cell embryo indicated.

(D) Pearson correlation coefficients for sister blastomeres of 2-cell embryos from wild-type, *Padi6*-C663A and *Padi6*-KO females. Unpaired two-tailed Student's t-test; *$P \leq 0.05$, **$P \leq 0.01$, ***$P \leq 0.001$, ****$P \leq 0.0001$, ns ($P > 0.05$).

(E) Transcript counts of *Padi1*, *Padi2*, *Padi3*, *Padi4* and *Padi6* in MII oocytes, zygotes and 2-cell embryos from wild-type, *Padi6*-C663A and *Padi6*-KO females.

(F) Venn diagrams depicting overlap of down and upregulated genes in zygotes and 2-cell embryos from *Padi6*-C663A and *Padi6*-KO females.

(G) GO analysis most enriched biological process (BP) terms for *Padi6*-C663A and *Padi6*-KO 2-cell embryo upregulated and downregulated genes respectively. GO analysis was performed using the Database for Annotation, Visualization and Integrated Discovery (DAVID)[90,91]. p-values were adjusted with the Benjamini-Hochberg procedure.

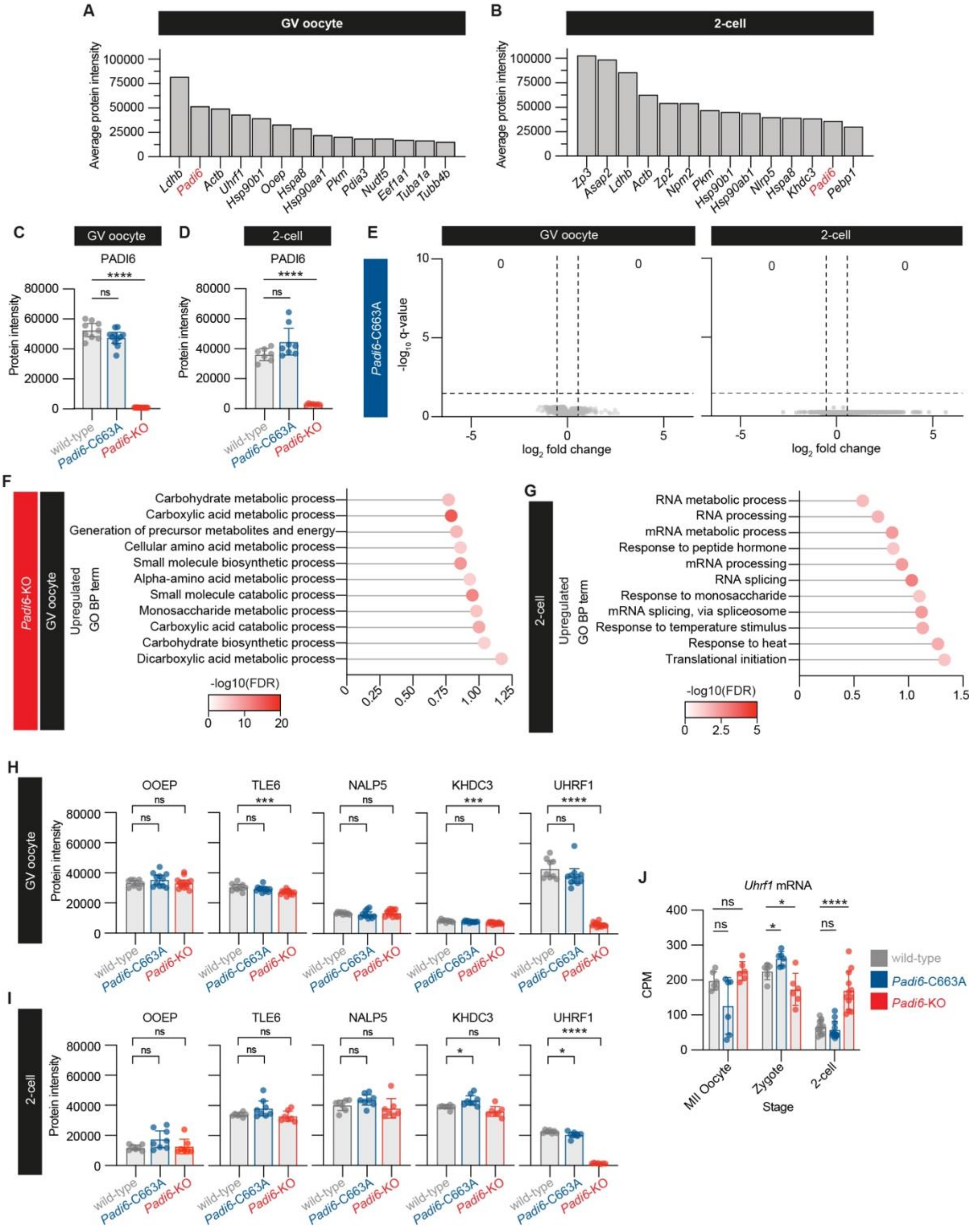


**Figure S5. Proteomic analysis of GV oocytes and 2-cell embryos from wild-type, *Padi6*$^{C663A/C663A}$ and *Padi6*$^{-/-}$ females. Related to Figure 4.**

(A and B) Average protein intensity values for the top 15 most abundant proteins identified in wild-type GV oocytes and 2-cell embryos.

(C and D) Protein intensity values of PADI6 in GV oocytes and 2-cell embryos from wild-type (GV oocyte *n* = 9, 2-cell *n* = 7), *Padi6*$^{C663A/C663A}$ (*Padi6*-C663A, GV oocyte *n* = 9, 2-cell *n* = 7) and *Padi6*$^{-/-}$ (*Padi6*-KO, GV oocyte *n* = 13, 2-cell *n* = 7) females. Unpaired two-tailed Student's t-test; **P* ≤ 0.05, ***P* ≤ 0.01, ****P* ≤ 0.001, *****P* ≤ 0.0001, ns (*P* > 0.05).

(E) Volcano plot of differentially abundant proteins in GV oocytes and 2-cell embryos from *Padi6*-C663A females compared with those from wild-type females. Dashed lines indicates q-value = 0.05 and $log_2$ fold change = -0.5 and 0.5.
(F) Enriched GO BP terms after STRING[59] analysis of proteins upregulated in GV oocytes from *Padi6*-KO females. Dot colour indicates $-log_{10}$ FDR. GO BP terms were filtered prior to plotting to include only those with an enrichment strength > 0.75 and FDR < 0.00005.
(G) Enriched GO BP terms after STRING[59] analysis of proteins upregulated in 2-cell embryos from *Padi6*-KO females. Dot colour indicates $-log_{10}$ FDR. GO BP terms were filtered prior to plotting to include only those with an enrichment strength > 0.5.
(H and I) Protein intensity values of OOEP, TLE6, KHDC3, NALP5 and UHRF1 in GV oocytes and 2-cell embryos from wild-type (GV oocyte *n* = 9, 2-cell *n* = 7), $Padi6^{C663A/C663A}$ (*Padi6*-C663A, GV oocyte *n* = 9, 2-cell *n* = 7) and $Padi6^{-/-}$ (*Padi6*-KO, GV oocyte *n* = 13, 2-cell *n* = 7) females. Unpaired two-tailed Student's t-test; *$P \leq 0.05$, **$P \leq 0.01$, ***$P \leq 0.001$, ****$P \leq 0.0001$, ns ($P > 0.05$).
(J) *Uhrf1* counts per million (CPM) in MII oocytes, zygotes, and 2-cell embryos from wild-type, *Padi6*-C663A and *Padi6*-KO females. Per genetic background MII oocytes (*n* = 6), zygotes (*n* = 6), 2-cell blastomeres (*n* = 12). Unpaired two-tailed Student's t-test with Holm-Šídák correction for multiple tests; *$P \leq 0.05$, **$P \leq 0.01$, ***$P \leq 0.001$, ****$P \leq 0.0001$, ns ($P > 0.05$).

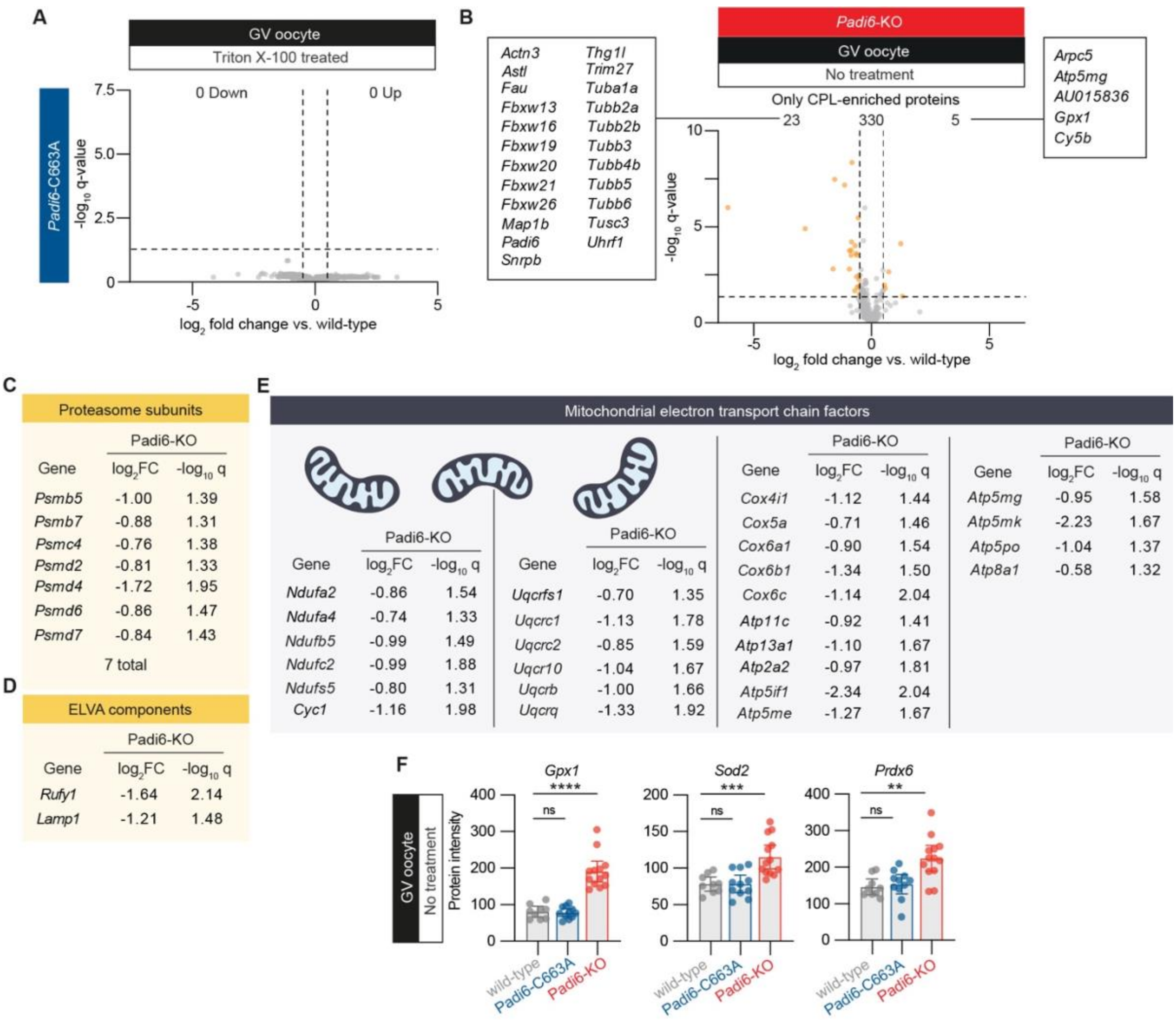


**Figure S6. Analysis of CPL-enriched proteins. Related to Figures 5 and 6.**

(A) Volcano plot of fold-changes of proteins in GV oocytes from *Padi6*$^{C663A/C663A}$ (*Padi6*-C663A) females compared with those from wild-type females. Dashed lines indicate q-value = 0.05 and $\log_2$ fold changes = -0.5 and 0.5.

(B) Volcano plot of differentially abundant proteins in GV oocytes from *Padi6*-KO females compared with those from wild-type females, filtered to only include proteins that were CPL-enriched. Orange points represent downregulated and upregulated proteins present with $\log_2$ fold change < -0.5 or > 0.5 respectively, and q-value < 0.05. Dashed lines indicate q-value = 0.05 and $\log_2$ fold change = -0.5 and 0.5.

(C) Proteasome subunits found to be enriched in the CPL fraction. The $\log_2$ fold changes and -$\log_{10}$ q-values of each protein calculated by comparing Triton X-100 treated *Padi6*$^{-/-}$ (*Padi6*-KO) GV oocytes compared with Triton X-100 treated wild-type GV oocytes are shown.

(D) ELVA proteins RUFY1 and LAMP1 found to be enriched in the CPL fraction. The $\log_2$ fold changes and -$\log_{10}$ q-values of each protein calculated by comparing Triton X-100 treated *Padi6*-KO GV oocytes compared with Triton X-100 treated wild-type GV oocytes are shown.

(E) Mitochondrial electron transport chain factors found to be CPL-enriched in the CPL fraction. The $\log_2$ fold changes and -$\log_{10}$ q-values of each protein calculated by comparing Triton X-100 treated *Padi6*-KO GV oocytes compared with Triton X-100 treated wild-type GV oocytes are shown.

(F) Protein intensity values of proteins upregulated during oxidative stress to counter ROS[93], in GV oocytes from wild-type ($n$ = 9), *Padi6*$^{C663A/C663A}$ (*Padi6*-C663A, $n$ = 9) and *Padi6*$^{-/-}$ (*Padi6*-KO, $n$ = 13) females. Unpaired two-tailed Student's t-test; *$P \leq 0.05$, **$P \leq 0.01$, ***$P \leq 0.001$, ****$P \leq 0.0001$, ns ($P > 0.05$).

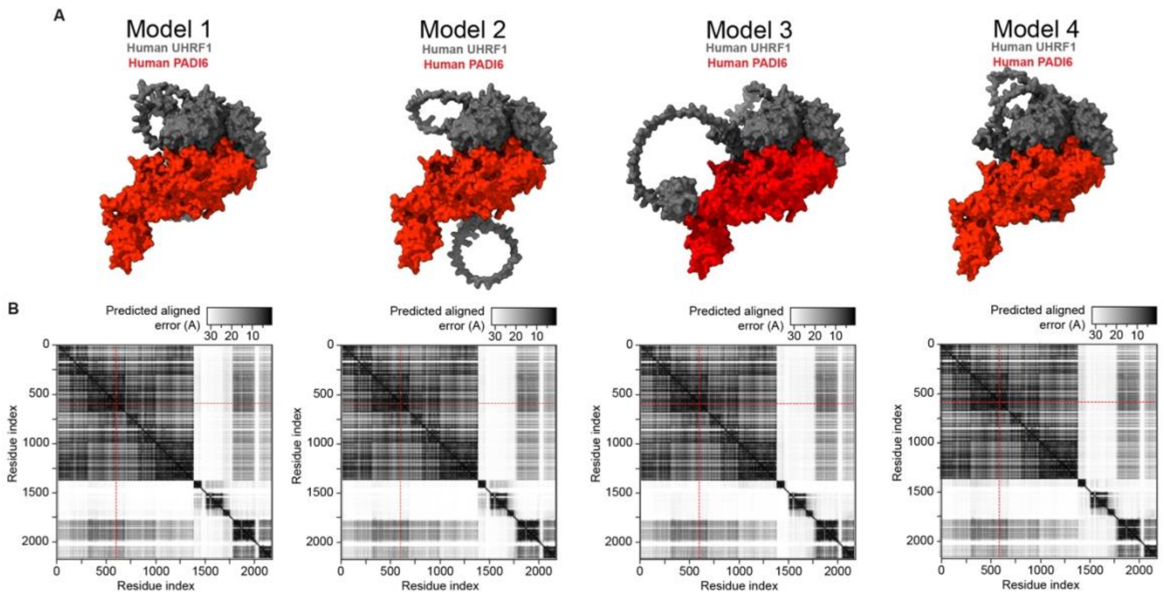


**Figure S7. AlphaFold Server predicted interaction between human UHRF1 and human PADI6. Related to Figure 6.**

(A) Models 1 to 4 of the AlphaFold Server[72,73] predicted interaction between hPADI6 (red) and UHRF1 (grey). Visualised with ChimeraX[74,75].

(B) Predicted aligned error heatmaps of models 1 to 4 of the AlphaFold predicted interaction between hPADI6 and UHRF1. hPADI6-N598 is highlighted by a red dashed line. Residue indices: hPADI6 molecule 1: 0-693, hPADI6 molecule 2: 694-1388, UHRF1 molecule: 1389-2182.

**Table S1. Padi6 C663A generation guide RNA sequences**

| Guide name | Guide Sequence (5'-) | Predicted cut site |
|---|---|---|
| Padi6 sgRNA-1 | GACTGCTACCTGGCCAACAT | chr4:140,727,650 |
| Padi6 sgRNA-2 | CTGCTACCTGGCCAACATAG | chr4:140,727,649 |
| Padi6 sgRNA-3 | CACCCTGTTTATGATGGCAC | chr4:140,727,628 |

**Table S2. Padi6 C663A ssODN sequence**

| | |
|---|---|
| Padi6 C663A ssODN | gtctcaagtgcaccttcattgatgattttgactgctacctggccaatataggcgacgtcGCCgctagtgccatcataaa cagggtgccatttgcattcaagtggtggaagatgaccccat |

**Table S3. Primer Sequences for Padi6 C663A genotyping by MiSeq**

| Amplicon PCR primers | Sequence (5'-3') |
|---|---|
| Padi6 MiSeq Forward | CGTGTTGGGCAAGAACCTTG |
| Padi6 MiSeq Reverse | GCGCTCTTTTCCCTTTTCCC |

**Table S4. Primer Sequences for Padi6 C663A long read sequencing**

ONT = Oxford Nanopore Technologies

| Long-read sequencing primers | Sequence (5'-3') |
|---|---|
| Padi6 ONT Forward | ACTCCACCTTAGGCCTGCTA |
| Padi6 ONT Reverse | GCCAGTGTTGCTCCAAGTTG |

**Table S5. Primer Sequences for Padi6 C663A ddPCR**

ddPCR = droplet digital polymerase chain reaction

| ddPCR primers | Sequence (5'-3') |
|---|---|
| Padi6 Mut ddPCR Reverse | GCAAATGGCACCCTGTTTATGATG |
| Padi6 Mut ddPCR Probe | TCGCCGCTAGTGCCATCATAAACA |

**Table S6. In-Fusion cloning primers**

| Product | Primer | Sequence (5'-3') |
|---|---|---|
| pcDNA3.1_Strep-Strep-TEV-mPADI6 | Vector Forward | TAATAATAATCTAGAGGGCCCGTTTAAACCC |
| | Vector Reverse | GCTGCCGCTACCTGACT |
| | Insert Forward | TCAGGTAGCGGCAGCATGTCTTTTCAGAACTCACTCAGCC |

| | | |
|---|---|---|
| | Insert Reverse | CCTCTAGATTATTATTATGGGGTCATCTTCCACCACT |
| pCMV_3XFlag-hPADI6 | Vector Forward | CCAGTCGACTCTAGAGGATCCCG |
| | Vector Reverse | TATCGATGAATTCGCGGCCG |
| | Insert Forward | GCGAATTCATCGATAATGGTCAGCGTGGAGGG |
| | Insert Reverse | TCTAGAGTCGACTGGCTAAGGTACCATCTTCCACCATTTGAAG |
| pCMV_HA-UHRF1 | Vector Forward | TGAGCGGCCGCGGGGATC |
| | Vector Reverse | GAATTCCTCCATGGCCATAAGAGC |
| | Insert Forward | GCCATGGAGGAATTCATGTGGATCCAGGTTCGGAC |
| | Insert Reverse | CCCCGCGGCCGCTCACTACCGGCCATTGCCG |
| pDNA3-MYC-Cullin1 | Insert Forward | CAGAGGAGGACCTGGAATTCATGTCGTCAACCCGGAGC |
| | Insert Reverse | GATGCATGCTCGAGCGGCCGCTTAAGCCAAGTAACTGTAGGTGTCCTTTTCACCA |
| pCDNA-MYC-UBE2D2 | Insert Forward | CAGAGGAGGACCTGGAATTCATGGCTCTGAAGAGAATCCACAAGGAATTGAATGA |
| | Insert Reverse | GATGCATGCTCGAGCGGCCGCTTACATCGCATACTTCTGAGTCCATTCCCGAGC |

**Table S7. Mutagenesis primers**

| **Mutation** | **Primer** | **Sequence (5’-3’)** |
|---|---|---|
| mPADI6$^{C663A}$ | Forward | CATAGGGGACGTCGCTGCCAGTGCCATCATAAAC |
| | Reverse | GTTTATGATGGCACTGGCAGCGACGTCCCCTATG |
| hPADI6$^{G540R}$ | Forward | GCAGATCAGCTCCTGTCTAATCGGAGGGAAGCCAAAACCATCG |
| | Reverse | CGATGGTTTTGGCTTCCCTCCGATTAGACAGGAGCTGATCTGC |
| hPADI6$^{N598S}$ | Forward | GTTCTGCTTGGAGAAGCTGACTAGCATCCCCTCTGACCAGCAG |
| | Reverse | CTGCTGGTCAGAGGGGATGCTAGTCAGCTTCTCCAAGCAGAAC |

**Table S8. Single oocyte/embryo DIA PASEF data acquisition windows**

| #MS Type | Cycle Id | Start IM [1/K0] | End IM [1/K0] | Start Mass [m/z] | End Mass [m/z] |
|---|---|---|---|---|---|
| MS1 | 0 | - | - | - | - |
| PASEF | 1 | 0.7 | 0.83 | 300.52 | 400.73 |
| PASEF | 1 | 0.83 | 1.3 | 604.84 | 620.02 |
| PASEF | 2 | 0.7 | 0.87 | 400.73 | 426.24 |
| PASEF | 2 | 0.87 | 1.3 | 620.02 | 635.64 |
| PASEF | 3 | 0.7 | 0.88 | 426.24 | 444.26 |
| PASEF | 3 | 0.88 | 1.3 | 635.64 | 652.36 |
| PASEF | 4 | 0.7 | 0.88 | 444.26 | 459.71 |
| PASEF | 4 | 0.88 | 1.3 | 652.36 | 670.36 |
| PASEF | 5 | 0.7 | 0.89 | 459.71 | 473.77 |
| PASEF | 5 | 0.89 | 1.3 | 670.36 | 689.85 |
| PASEF | 6 | 0.7 | 0.9 | 473.77 | 487.74 |
| PASEF | 6 | 0.9 | 1.3 | 689.85 | 710.34 |
| PASEF | 7 | 0.7 | 0.91 | 487.74 | 501.24 |
| PASEF | 7 | 0.91 | 1.3 | 710.34 | 732 |
| PASEF | 8 | 0.7 | 0.92 | 501.24 | 514.27 |
| PASEF | 8 | 0.92 | 1.3 | 732 | 756.89 |
| PASEF | 9 | 0.7 | 0.93 | 514.27 | 526.74 |
| PASEF | 9 | 0.93 | 1.3 | 756.89 | 783.41 |
| PASEF | 10 | 0.7 | 0.93 | 526.74 | 539.29 |
| PASEF | 10 | 0.93 | 1.3 | 783.41 | 813.35 |
| PASEF | 11 | 0.7 | 0.94 | 539.29 | 552.24 |
| PASEF | 11 | 0.94 | 1.3 | 813.35 | 845.89 |
| PASEF | 12 | 0.7 | 0.95 | 552.24 | 564.79 |
| PASEF | 12 | 0.95 | 1.3 | 845.89 | 884.43 |
| PASEF | 13 | 0.7 | 0.97 | 564.79 | 577.61 |
| PASEF | 13 | 0.97 | 1.3 | 884.43 | 934.5 |
| PASEF | 14 | 0.7 | 0.98 | 577.61 | 590.95 |
| PASEF | 14 | 0.98 | 1.3 | 934.5 | 1009.48 |
| PASEF | 15 | 0.7 | 1.02 | 590.95 | 604.84 |
| PASEF | 15 | 1.02 | 1.3 | 1009.48 | 1198.99 |

**Figure S8: Uncut western blot images with replicates for Figure 6C**

Dashed square boxes indicate regions used in figure.

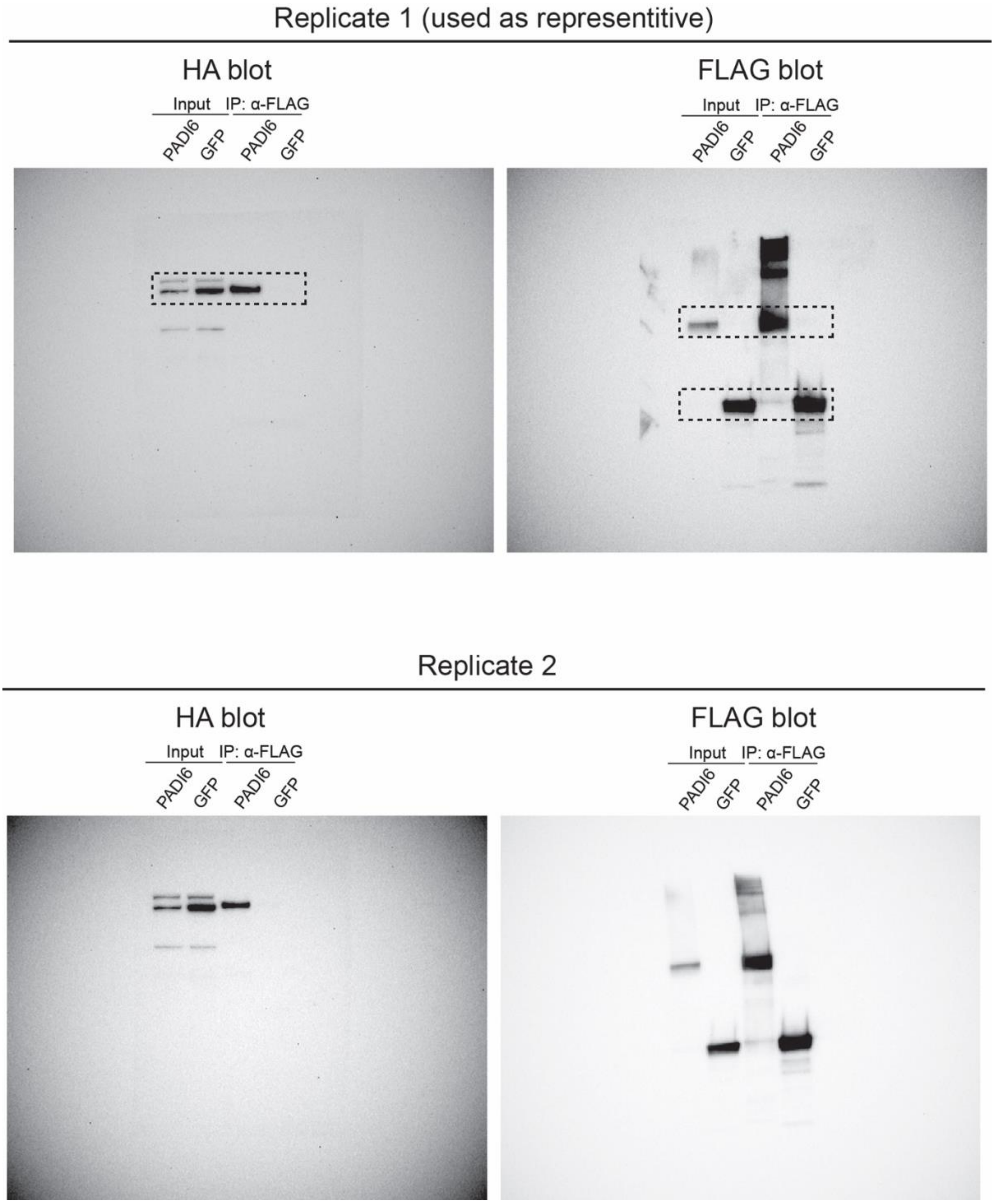

**Figure S9: Uncut western blot images with replicates for Figure 6G**

Dashed square boxes indicate regions used in figure.

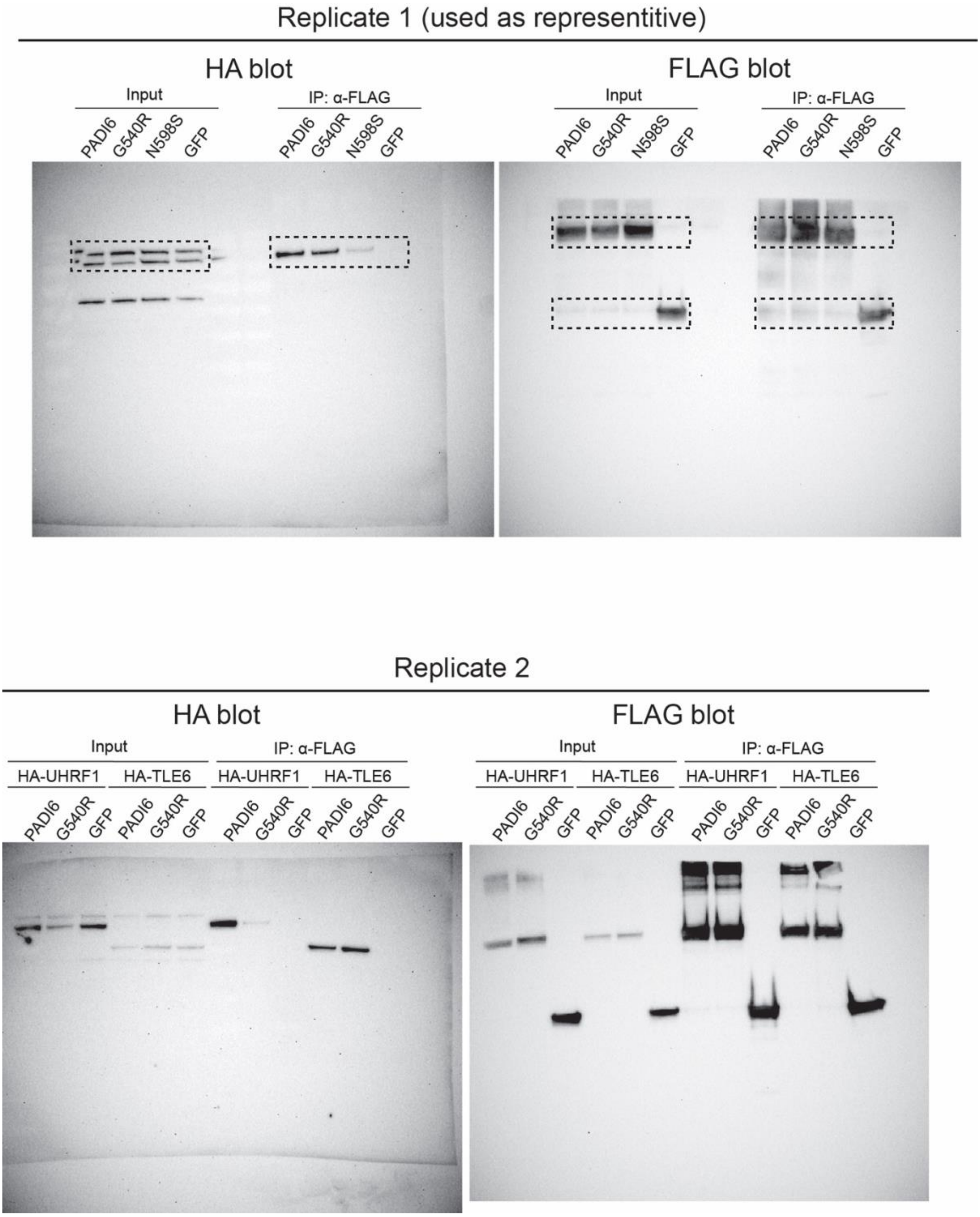

**Figure S10: Uncut western blot images with replicates for Figure 6I**

Dashed square boxes indicate regions used in figure.

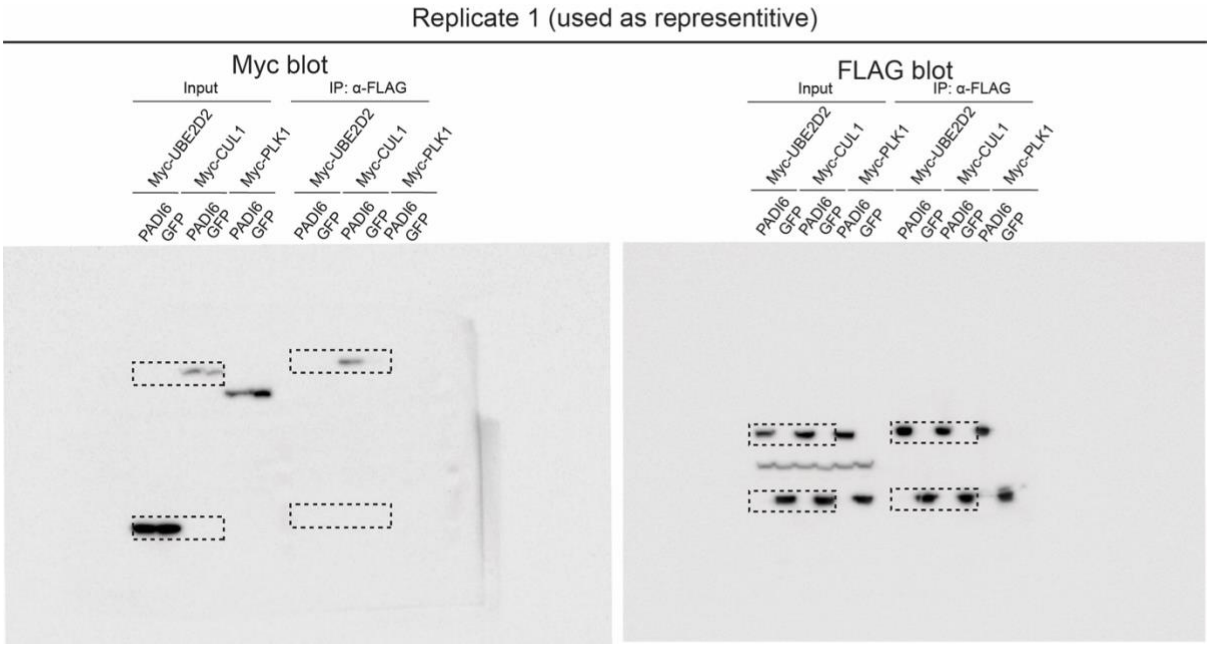

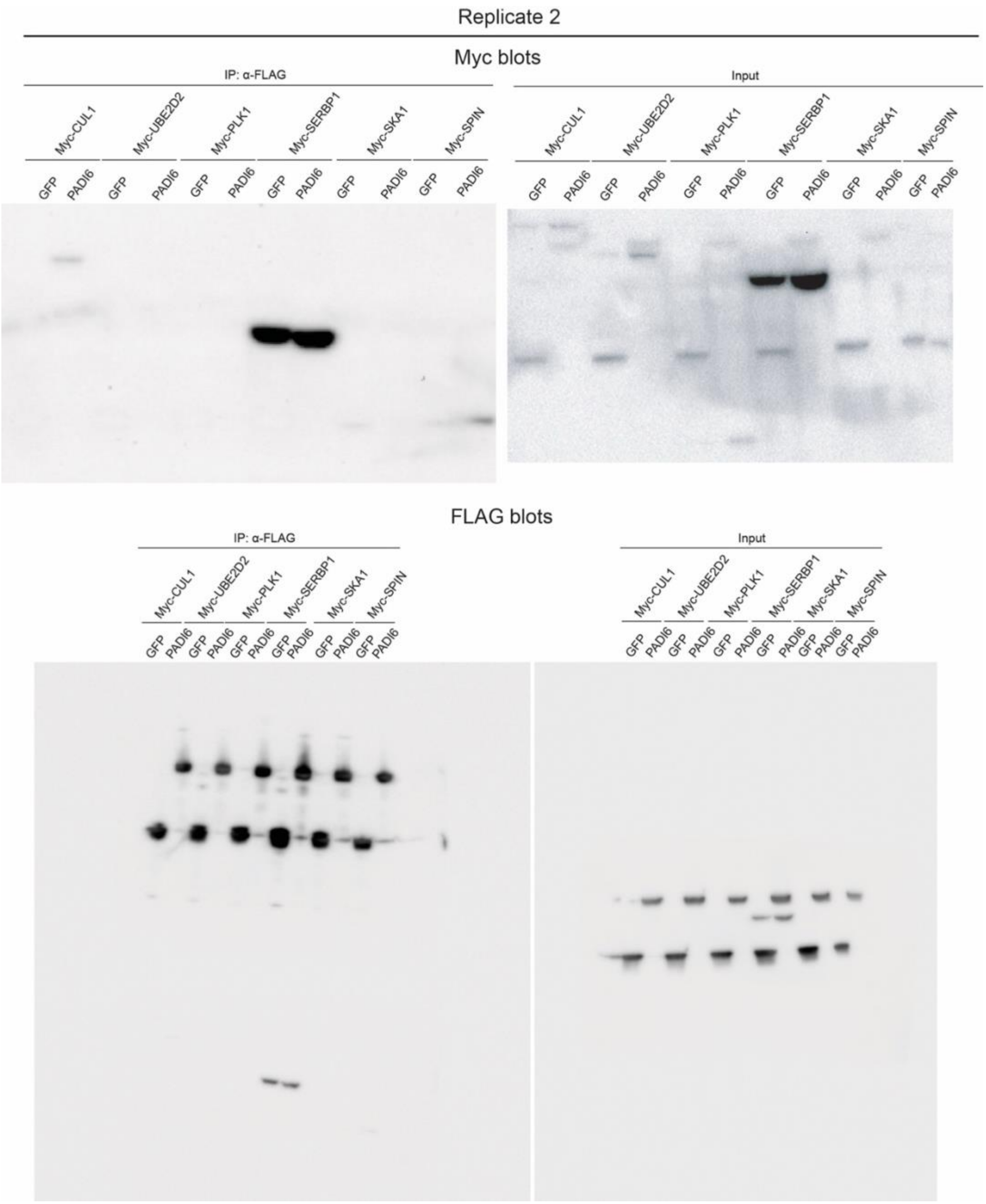
Replicate 2
Myc blots
IP: α-FLAG
Input
Myc-CUL1
Myc-UBE2D2
Myc-PLK1
Myc-SERBP1
Myc-SKA1
Myc-SPIN
GFP
PADI6
FLAG blots
